# Exploring the tymovirids landscape through metatranscriptomics data

**DOI:** 10.1101/2021.07.15.452586

**Authors:** Nicolás Bejerman, Humberto Debat

## Abstract

*Tymovirales* is an order of viruses with positive-sense, single-stranded RNA genomes that mostly infect plants, but also fungi and insects. The number of tymovirid sequences has been growing in the last few years with the extensive use of high-throughput sequencing platforms. Here we report the discovery of 31 novel tymovirid genomes associated with 27 different host plant species, which were hidden in public databases. These viral sequences were identified through a homology searches in more than 3,000 plant transcriptomes from the NCBI Sequence Read Archive (SRA) using known tymovirids sequences as query. Identification, assembly and curation of raw SRA reads resulted in 29 viral genome sequences with full-length coding regions, and two partial genomes. Highlights of the obtained sequences include viruses with unique and novel genome organizations among known tymovirids. Phylogenetic analysis showed that six of the novel viruses were related to alphaflexiviruses, seventeen to betaflexiviruses, two to deltaflexiviruses and six to tymoviruses. These findings resulted in the most complete phylogeny of tymovirids to date and shed new light on the phylogenetic relationships and evolutionary landscape of this group of viruses. Furthermore, this study illustrates the complexity and diversity of tymovirids genomes and demonstrates that analyzing SRA public data provides an invaluable tool to accelerate virus discovery and refine virus taxonomy.

## Introduction

The application of high-throughput sequencing (HTS) is becoming a fundamental tool in biomedical research, resulting in a steep increase in the number of genome and transcriptome sequencing projects that is leading to a massive number of nucleotides deposited in the Sequence Read Archive (SRA) of the National Center for Biotechnology Information (NCBI). Over 16,000 petabases (10^15^ bases) have been deposited in the SRA, with over 6,000 petabases available as open-access data [1], and this number is increasing significantly each year. This huge amount of data provides a significant challenge derived in the high costs of data storage and management. Given the nature, objectives and magnitude of the sequencing projects most of the generated data remains largely unexplored.

In the last years the application of metagenomics approaches resulted in the discovery of an abundant number of novel viruses. Most of them likely do not induce any apparent symptoms in their host or their host is unknown. In addition, the detection of multiple viruses in single samples suggest that mixed infections are common and represent an important constrain to an unbiased assessment of the contribution of each virus in a tentative pathosystem. Importantly, detections in metagenomics datasets usually correspond to novel and unreported viruses. This illustrates how limited is our knowledge about the richness of the plant virosphere, that seems to be exceedingly diverse in every host assessed so far [2–4]. Even though the application of HTS has resulted in the discovery of a great number of viruses, the pace of discovery suggest that it is still a minuscule portion of the whole virosphere, which has led to initiatives to accommodate this massive diversity in a proposal for a first comprehensive megataxonomy of the virus world [5].

The submitters of transcriptome datasets often limit their scientific interest to their specific field of study, leaving behind a large amount of unused data which is potentially valuable [6]. Several studies in the last few years have been focusing in the analysis of such transcriptome datasets, where viral sequences could be hidden in plain sight, which resulted in the discovery of novel viral sequences [7–13]. Simmonds and colleagues [14] consensually stated that viruses that are known only from metagenomic data can, should, and have been incorporated into the official classification scheme overseen by the International Committee on Taxonomy of Viruses (ICTV). Therefore, analyzing public sequence databases is becoming a great source for mining novel plant viruses, that results in the identification of novel viruses in hosts with no previous record of virus infections [7]. The analysis of publically available databases does not require the acquisition of samples and subsequent sequencing, resulting in an inexpensive and sustainable approach for virus discovery. Proper attribution to the generators of the original data is essential to promote the good practices of open and transparent research, which enable the production of new results with existing data. Thus, the secondary analysis of publically available transcriptomic data to address novel research question and objectives is likely a wide-ranging, efficient and cost-effective approach for virus discovery due to the millions of datasets from a vast variety of potential host species available at the NCBI-SRA [11].

Tymovirales is an order of viruses which encompass viruses with a positive-sense, single-stranded RNA genome that mostly infect plants, but some are associated to fungi, while others are associated with insects. This order is taxonomically classified into five families: *Alphaflexiviridae, Betaflexiviridae, Deltaflexiviridae, Gammaflexiviridae*, whose members have flexuous filamentous virions, and *Tymoviridae*, whose members have isometric virions [15].

The family *Alphaflexiviridae* contains seven genera: *Allexivirus Lolavirus, Mandarivirus, Platypuvirus* and *Potexvirus* genera comprised by plant-associated viruses, while *Botrexvirus* and *Sclerodarnavirus* genera correspond to fungi-associated viruses [16]. However, a recent proposal suggested to abolish the genus *Mandarivirus* and create the subgenus *Mandarivirus* within the genus *Potexvirus* [17].

Alphaflexiviruses have a polyadenylated genome of 5.9-9.0 kb in size with 1 or 5 open reading frames (ORFs) in those fungi-associated viruses and 5 to 7 ORFs in those plant-infecting viruses. The gene encoding the replicase (REP) is the only conserved among all alphaflexivirues. In plant-infecting alphafleviruses, the gene encoding the capsid protein (CP) and, with the exception of platypuviruses, the module of the three overlapping ORFs 2 to 4, which encode the triple gene block (TGB) that is involved in the cell-to-cell movement, are conserved [16].

The family *Betaflexiviridae* is composed by plant-infecting viruses, classified into two subfamilies, *Quinvirinae* and *Trivirinae*, distinguished by the cell-to-cell type movement protein (MP), 30K-like or TGB, that are classified in thirteen genera. *Carlavirus, Foveavirus*, and *Robigovirus* genera belong to the *Quinvirianae* subfamily, while *Capillovirus, Chordovirus, Citrivirus, Divavirus, Prunevirus, Ravavirus, Tepovirus, Trichivirus, Tepovirus*, and *Wamavirus* genera belong to the *Trinvirinae* subfamily [15].

Betaflexiviruses have a polyadenylated genome of 6.5-9.5 kb in size with 2 to 6 ORFs. Each betaflexivirus typically codes for a REP, a MP, and a CP; while some viruses also code for a nucleic acid binding protein (NABP) [15]. The viruses belonging to the subfamily *Quinvirinae* have 5 to 6 ORFs, while those viruses in the subfamily *Trivirinae* have 2 to 5 ORFs [15]. Among viruses belonging to the later subfamily there is variation in their genome organization. For instance, the MP encoded by capilloviruses and divaviruses is nested within the ORF1, while that one of citriviruses and trichoviruses overlap with the REP [15].

The family *Deltaflexiviridae* contains one genera, named as *Deltaflexivirus*, which is comprised by fungi-associated viruses [18–20]. Deltaflexiviruses have a polyadenylated genome of 8.1-8.3 kb in size with 4 to 5 ORFs. The gene encoding the replicase is the only one conserved among all deltaflexiviruses [18–20]. The family *Tymoviridae* contains three genera the *Marafivirus* and the *Maculavirus*, which have a 3’ terminal poly (A) tail, and *Tymovirus*, whose genomes are not polyadenylated [21]. The genus *Marafivirus* is exclusively composed of plant-infecting viruses, but several tymoviruses have been associated to fungi or insects [22–24]. Tymoviruses have a genome of 6.0-7.5 kb in size with 1 to 4 ORFs, and the gene encoding the replicase is the only conserved among all tymoviruses [21].

In this study we queried the publically available plant transcriptome datasets in the transcriptome shotgun assembly (TSA) database and their corresponding SRA datasets hosted at NCBI and identified 31 novel tymovirids from 27 plant species, showing structural, functional and evolutionary cues to be classified in new and existing genera within the families *Alphaflexiviridae, Betaflexiviridae, Deltaflexiviridae* and *Tymoviridae* in the order *Tymovirales*. The discovery of these new viruses increase the knowledge of tymovirids and greatly expands the number of viruses associated to the order *Tymovirales*.

## Materials and methods

### Identification of tymovirid sequences from public plant transcriptome datasets

The amino acid sequences corresponding to the replicase proteins of several known viruses belonging to the order *Tymovirales*, such as Cnidium virus X (QVW10166), Fusarium graminearum deltaflexivirus 1 (ANS13830), alfalfa virus F (AWB13379), eggplant mosaic virus (P20126), apple stem grooving virus (QVX32671), Diuris virus A (AFV57238), fig fleck-associated virus 2 (AOF41056), Helenium virus S (QQX32728), and garlic yellow mosaic-associated virus (AZM69107), were used as query in tBlastn searches with parameters word size = 6, expected threshold = 10, and scoring matrix = BLOSUM62, against the viridiplantae (taxid:33090) TSA database. The obtained hits were explored by hand and based on percentage identity (<90%), query coverage (>50%) and E-value (>1e-5), shortlisted as likely corresponding to novel virus transcripts, which were further analyzed. Given the redundant nature of many retrieved hits, a step of contig clustering was implemented using the CD-hit suite with standard parameters available at http://weizhongli-lab.org/cdhit_suite/cgi-bin/index.cgi?cmd=cd-hit. In addition, the raw data corresponding to the SRA experiments associated with the different target NCBI Bioprojects (Table 1) was retrieved for further analyses.

**Table 1.**
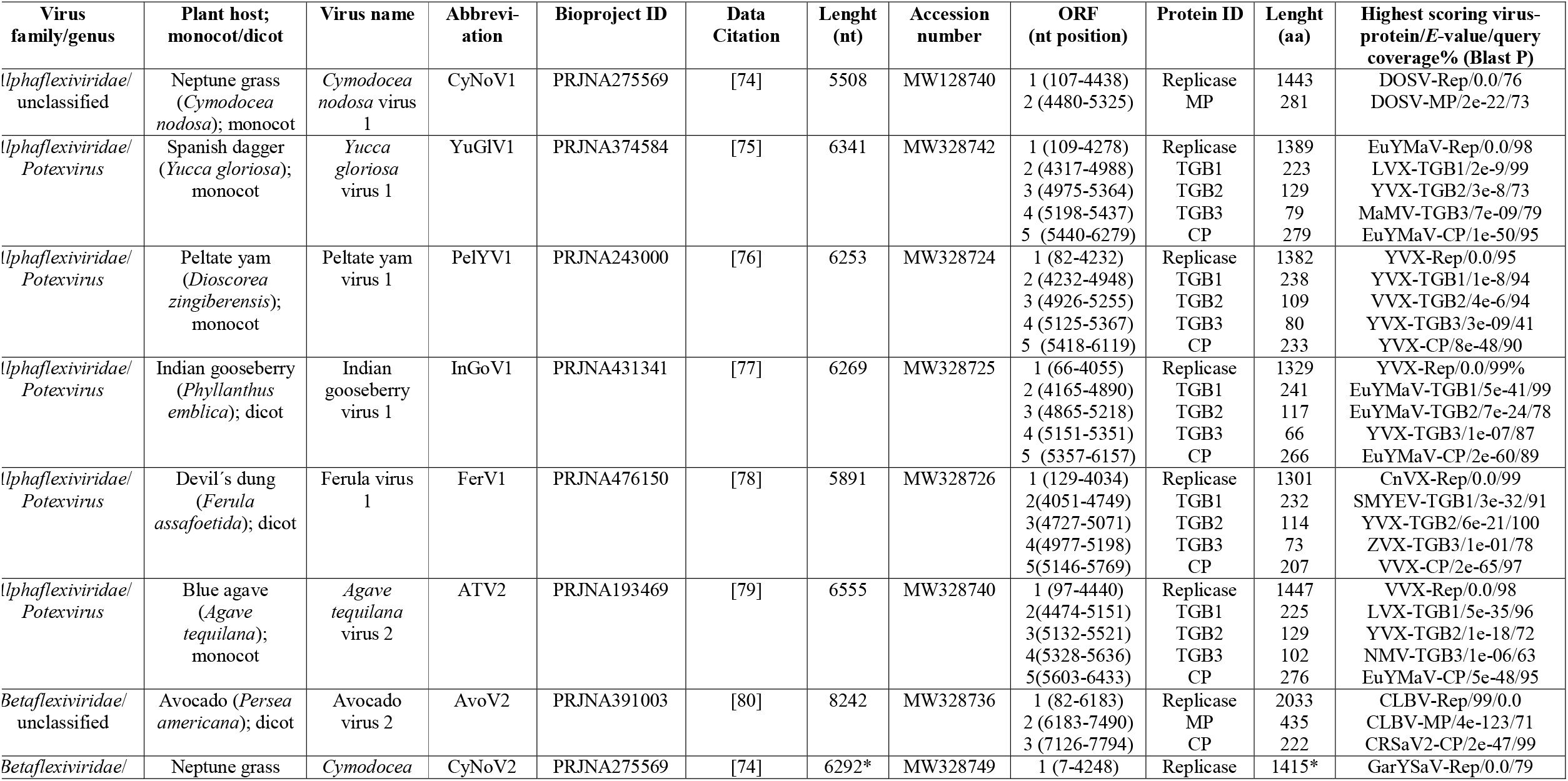
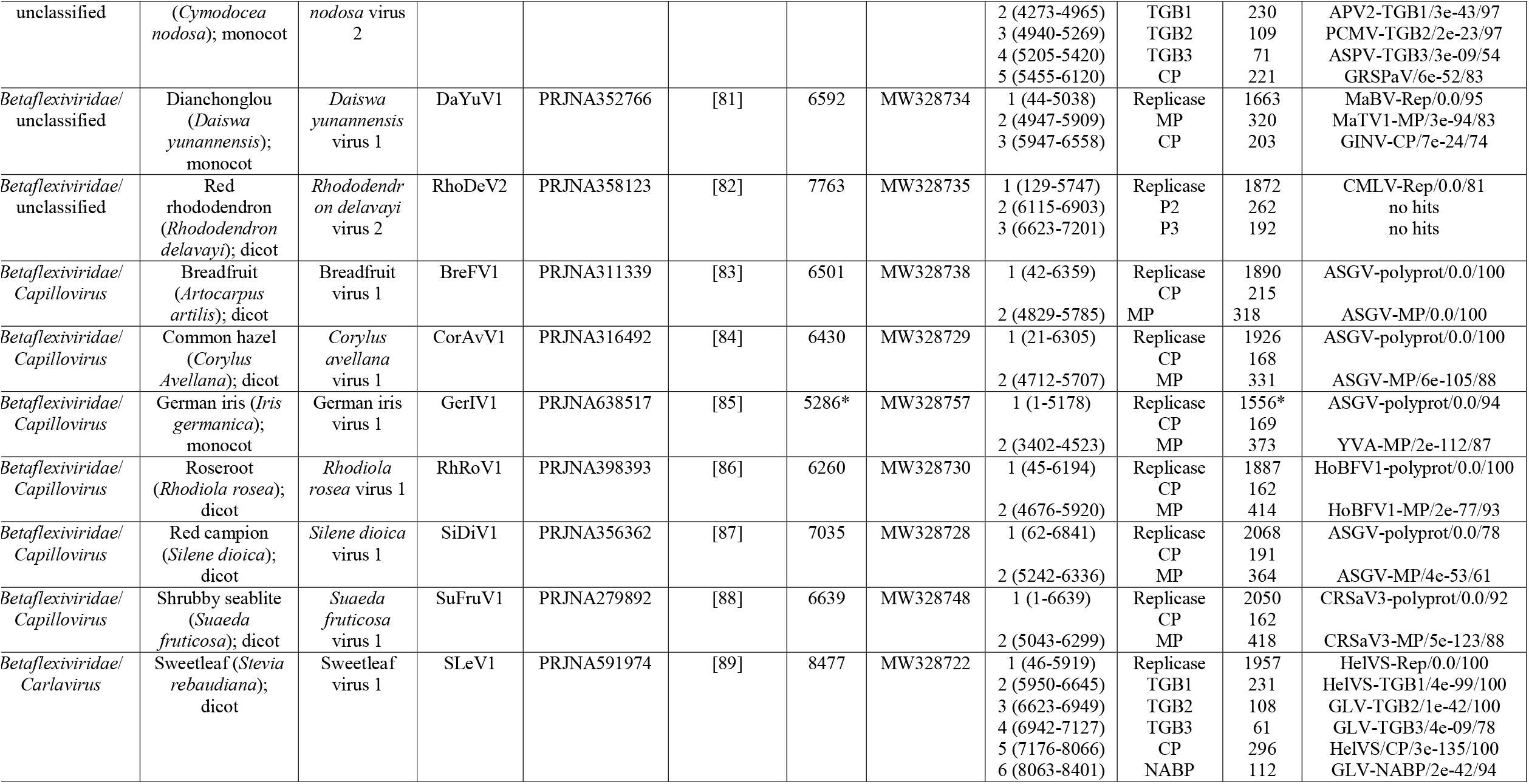
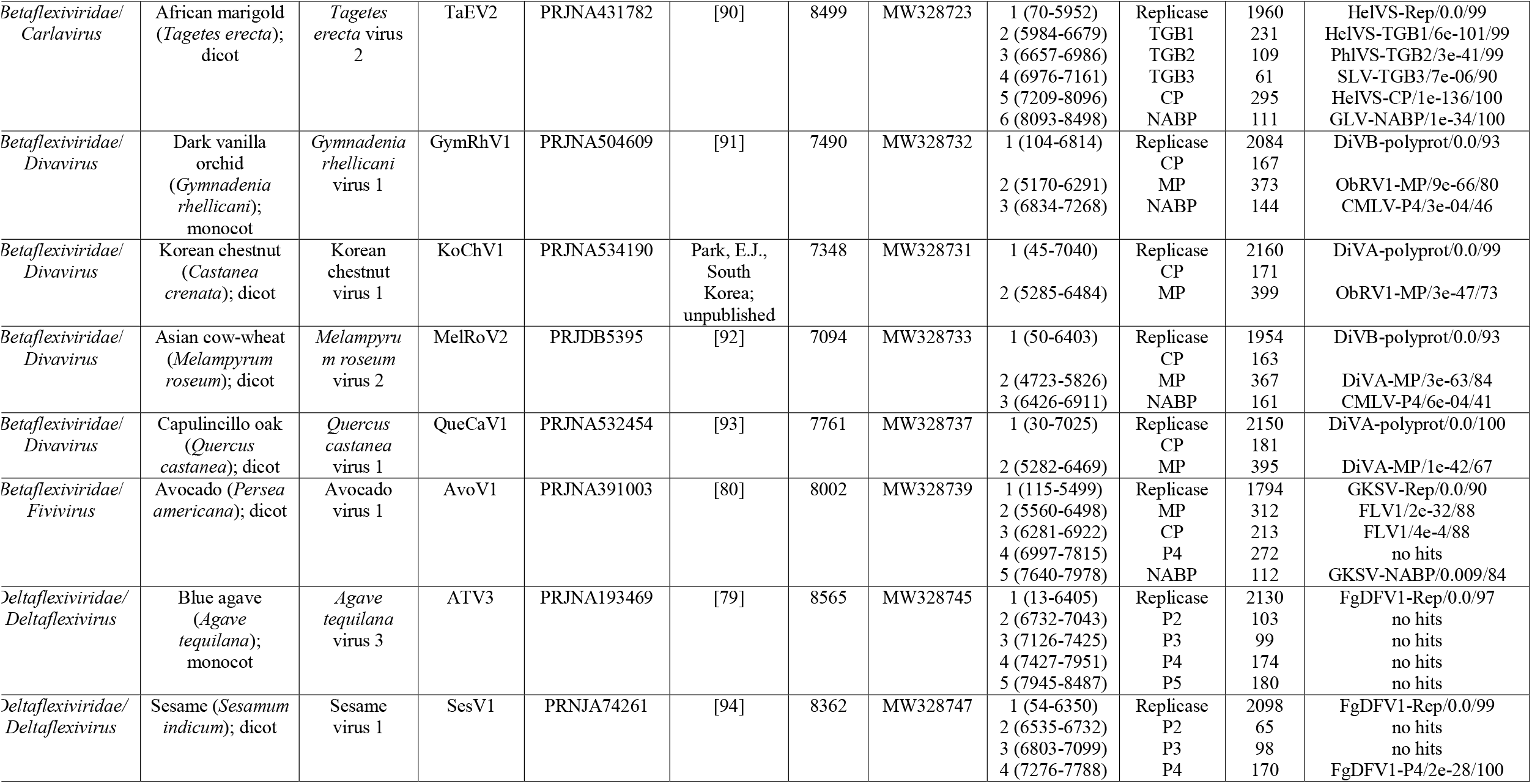
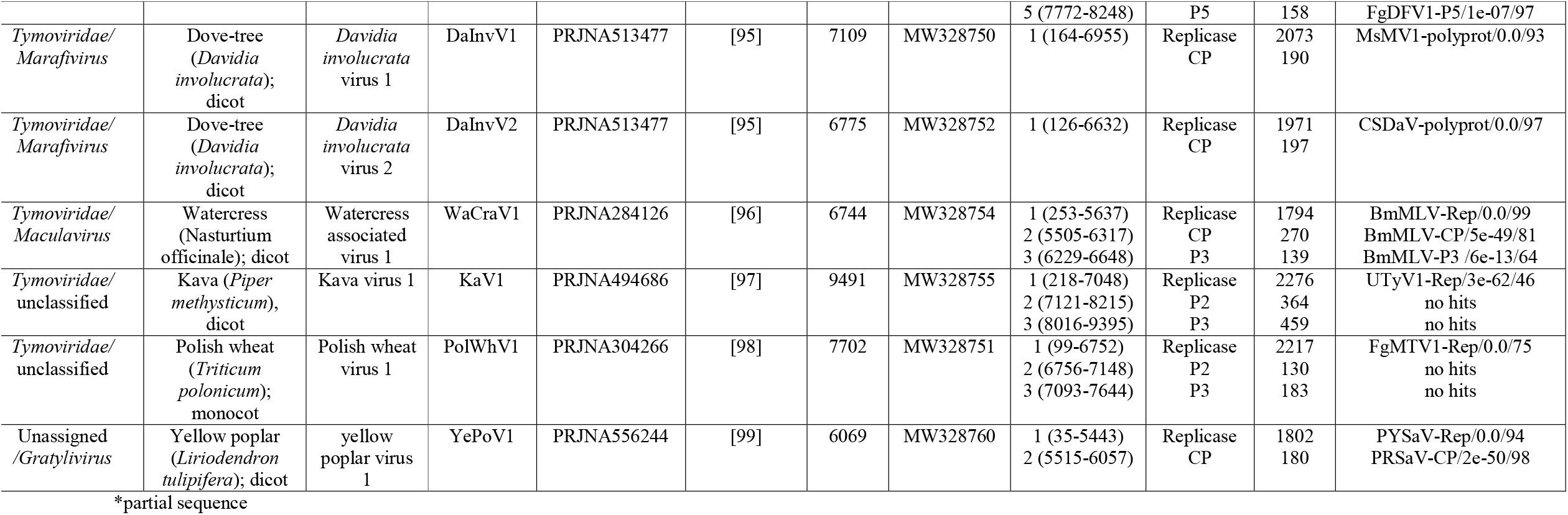
Summary of the tymovirids contigs identified from plant transcriptome data available in the NCBI database

| **Virus family/genus** | **Plant host; monocot/dicot** | **Virus name** | **Abbreviation** | **Bioproject ID** | **Data Citation** | **Lenght (nt)** | **Accession number** | **ORF (nt position)** | **Protein ID** | **Lenght (aa)** | **Highest scoring virus-protein/E-value/query coverage% (Blast P)** |
| --- | --- | --- | --- | --- | --- | --- | --- | --- | --- | --- | --- |
| *Alphaflexiviridae/*<br>unclassified | Neptune grass<br>(*Cymodocea nodosa*); monocot | *Cymodocea nodosa* virus 1 | CyNoV1 | PRJNA275569 | [74] | 5508 | MW128740 | 1 (107-4438)<br>2 (4480-5325) | Replicase<br>MP | 1443<br>281 | DOSV-Rep/0.0/76<br>DOSV-MP/2e-22/73 |
| *Alphaflexiviridae/*<br>*Potexvirus* | Spanish dagger<br>(*Yucca gloriosa*); monocot | *Yucca gloriosa* virus 1 | YuGIV1 | PRJNA374584 | [75] | 6341 | MW328742 | 1 (109-4278)<br>2 (4317-4988)<br>3 (4975-5364)<br>4 (5198-5437)<br>5 (5440-6279) | Replicase<br>TGB1<br>TGB2<br>TGB3<br>CP | 1389<br>223<br>129<br>79<br>279 | EuYMaV-Rep/0.0/98<br>LVX-TGB1/2e-9/99<br>YVX-TGB2/3e-8/73<br>MaMV-TGB3/7e-09/79<br>EuYMaV-CP/1e-50/95 |
| *Alphaflexiviridae/*<br>*Potexvirus* | Peltate yam<br>(*Dioscorea zingiberensis*); monocot | Peltate yam virus 1 | PeLYV1 | PRJNA243000 | [76] | 6253 | MW328724 | 1 (82-4232)<br>2 (4232-4948)<br>3 (4926-5255)<br>4 (5125-5367)<br>5 (5418-6119) | Replicase<br>TGB1<br>TGB2<br>TGB3<br>CP | 1382<br>238<br>109<br>80<br>233 | YVX-Rep/0.0/95<br>YVX-TGB1/1e-8/94<br>VVX-TGB2/4e-6/94<br>YVX-TGB3/3e-09/41<br>YVX-CP/8e-48/90 |
| *Alphaflexiviridae/*<br>*Potexvirus* | Indian gooseberry<br>(*Phyllanthus emblica*); dicot | Indian gooseberry virus 1 | InGoV1 | PRJNA431341 | [77] | 6269 | MW328725 | 1 (66-4055)<br>2 (4165-4890)<br>3 (4865-5218)<br>4 (5151-5351)<br>5 (5357-6157) | Replicase<br>TGB1<br>TGB2<br>TGB3<br>CP | 1329<br>241<br>117<br>66<br>266 | YVX-Rep/0.0/99%<br>EuYMaV-TGB1/5e-41/99<br>EuYMaV-TGB2/7e-24/78<br>YVX-TGB3/1e-07/87<br>EuYMaV-CP/2e-60/89 |
| *Alphaflexiviridae/*<br>*Potexvirus* | Devil's dung<br>(*Ferula assafoetida*); dicot | Ferula virus 1 | FerV1 | PRJNA476150 | [78] | 5891 | MW328726 | 1 (129-4034)<br>2(4051-4749)<br>3(4727-5071)<br>4(4977-5198)<br>5(5146-5769) | Replicase<br>TGB1<br>TGB2<br>TGB3<br>CP | 1301<br>232<br>114<br>73<br>207 | CnVX-Rep/0.0/99<br>SMYEV-TGB1/3e-32/91<br>YVX-TGB2/6e-21/100<br>ZVX-TGB3/1e-01/78<br>VVX-CP/2e-65/97 |
| *Alphaflexiviridae/*<br>*Potexvirus* | Blue agave<br>(*Agave tequilana*); monocot | *Agave tequilana* virus 2 | ATV2 | PRJNA193469 | [79] | 6555 | MW328740 | 1 (97-4440)<br>2(4474-5151)<br>3(5132-5521)<br>4(5328-5636)<br>5(5603-6433) | Replicase<br>TGB1<br>TGB2<br>TGB3<br>CP | 1447<br>225<br>129<br>102<br>276 | VVX-Rep/0.0/98<br>LVX-TGB1/5e-35/96<br>YVX-TGB2/1e-18/72<br>NMV-TGB3/1e-06/63<br>EuYMaV-CP/5e-48/95 |
| *Betaflexiviridae/*<br>unclassified | Avocado (*Persea americana*); dicot | Avocado virus 2 | AvoV2 | PRJNA391003 | [80] | 8242 | MW328736 | 1 (82-6183)<br>2 (6183-7490)<br>3 (7126-7794) | Replicase<br>MP<br>CP | 2033<br>435<br>222 | CLBV-Rep/99/0.0<br>CLBV-MP/4e-123/71<br>CRSaV2-CP/2e-47/99 |
| *Betaflexiviridae/* | Neptune grass | *Cymodocea* | CyNoV2 | PRJNA275569 | [74] | 6292* | MW328749 | 1 (7-4248) | Replicase | 1415* | GarYSaV-Rep/0.0/79 |
| unclassified | (*Cymodocea nodosa*); monocot | *nodosa* virus 2 |  |  |  |  |  | 2 (4273-4965)<br>3 (4940-5269)<br>4 (5205-5420)<br>5 (5455-6120) | TGB1<br>TGB2<br>TGB3<br>CP | 230<br>109<br>71<br>221 | APV2-TGB1/3e-43/97<br>PCMV-TGB2/2e-23/97<br>ASPV-TGB3/3e-09/54<br>GRSPaV/6e-52/83 |
| *Betaflexiviridae*/<br>unclassified | Dianchonglou (*Daiswa yunannensis*); monocot | *Daiswa yunannensis* virus 1 | DaYuV1 | PRJNA352766 | [81] | 6592 | MW328734 | 1 (44-5038)<br>2 (4947-5909)<br>3 (5947-6558) | Replicase<br>MP<br>CP | 1663<br>320<br>203 | MaBV-Rep/0.0/95<br>MaTV1-MP/3e-94/83<br>GINV-CP/7e-24/74 |
| *Betaflexiviridae*/<br>unclassified | Red rhododendron (*Rhododendron delavayi*); dicot | *Rhododendron delavayi* virus 2 | RhoDeV2 | PRJNA358123 | [82] | 7763 | MW328735 | 1 (129-5747)<br>2 (6115-6903)<br>3 (6623-7201) | Replicase<br>P2<br>P3 | 1872<br>262<br>192 | CMLV-Rep/0.0/81<br>no hits<br>no hits |
| *Betaflexiviridae*/<br>*Capillovirus* | Breadfruit (*Artocarpus artilis*); dicot | Breadfruit virus 1 | BreFV1 | PRJNA311339 | [83] | 6501 | MW328738 | 1 (42-6359)<br>2 (4829-5785) | Replicase<br>CP<br>MP | 1890<br>215<br>318 | ASGV-polyprot/0.0/100<br>ASGV-MP/0.0/100 |
| *Betaflexiviridae*/<br>*Capillovirus* | Common hazel (*Corylus avellana*); dicot | *Corylus avellana* virus 1 | CorAvV1 | PRJNA316492 | [84] | 6430 | MW328729 | 1 (21-6305)<br>2 (4712-5707) | Replicase<br>CP<br>MP | 1926<br>168<br>331 | ASGV-polyprot/0.0/100<br>ASGV-MP/6e-105/88 |
| *Betaflexiviridae*/<br>*Capillovirus* | German iris (*Iris germanica*); monocot | German iris virus 1 | GerIV1 | PRJNA638517 | [85] | 5286* | MW328757 | 1 (1-5178)<br>2 (3402-4523) | Replicase<br>CP<br>MP | 1556*<br>169<br>373 | ASGV-polyprot/0.0/94<br>YVA-MP/2e-112/87 |
| *Betaflexiviridae*/<br>*Capillovirus* | Roseroot (*Rhodiola rosea*); dicot | *Rhodiola rosea* virus 1 | RhRoV1 | PRJNA398393 | [86] | 6260 | MW328730 | 1 (45-6194)<br>2 (4676-5920) | Replicase<br>CP<br>MP | 1887<br>162<br>414 | HoBFV1-polyprot/0.0/100<br>HoBFV1-MP/2e-77/93 |
| *Betaflexiviridae*/<br>*Capillovirus* | Red campion (*Silene dioica*); dicot | *Silene dioica* virus 1 | SiDiV1 | PRJNA356362 | [87] | 7035 | MW328728 | 1 (62-6841)<br>2 (5242-6336) | Replicase<br>CP<br>MP | 2068<br>191<br>364 | ASGV-polyprot/0.0/78<br>ASGV-MP/4e-53/61 |
| *Betaflexiviridae*/<br>*Capillovirus* | Shrubby seablite (*Suaeda fruticosa*); dicot | *Suaeda fruticosa* virus 1 | SuFruV1 | PRJNA279892 | [88] | 6639 | MW328748 | 1 (1-6639)<br>2 (5043-6299) | Replicase<br>CP<br>MP | 2050<br>162<br>418 | CRSaV3-polyprot/0.0/92<br>CRSaV3-MP/5e-123/88 |
| *Betaflexiviridae*/<br>*Carlavirus* | Sweetleaf (*Stevia rebaudiana*); dicot | Sweetleaf virus 1 | SLeV1 | PRJNA591974 | [89] | 8477 | MW328722 | 1 (46-5919)<br>2 (5950-6645)<br>3 (6623-6949)<br>4 (6942-7127)<br>5 (7176-8066)<br>6 (8063-8401) | Replicase<br>TGB1<br>TGB2<br>TGB3<br>CP<br>NABP | 1957<br>231<br>108<br>61<br>296<br>112 | HeIVS-Rep/0.0/100<br>HeIVS-TGB1/4e-99/100<br>GLV-TGB2/1e-42/100<br>GLV-TGB3/4e-09/78<br>HeIVS/CP/3e-135/100<br>GLV-NABP/2e-42/94 |
| *Betaflexiviridae/*<br>*Carlavirus* | African marigold<br>(*Tagetes erecta*);<br>dicot | *Tagetes*<br>*erecta* virus<br>2 | TaEV2 | PRJNA431782 | [90] | 8499 | MW328723 | 1 (70-5952)<br>2 (5984-6679)<br>3 (6657-6986)<br>4 (6976-7161)<br>5 (7209-8096)<br>6 (8093-8498) | Replicase<br>TGB1<br>TGB2<br>TGB3<br>CP<br>NABP | 1960<br>231<br>109<br>61<br>295<br>111 | HelVS-Rep/0.0/99<br>HelVS-TGB1/6e-101/99<br>PhlVS-TGB2/3e-41/99<br>SLV-TGB3/7e-06/90<br>HelVS-CP/1e-136/100<br>GLV-NABP/1e-34/100 |
| *Betaflexiviridae/*<br>*Divavirus* | Dark vanilla<br>orchid<br>(*Gymnadenia*<br>*rhellicani*);<br>monocot | *Gymnadenia*<br>*rhellicani*<br>virus 1 | GymRhV1 | PRJNA504609 | [91] | 7490 | MW328732 | 1 (104-6814)<br>2 (5170-6291)<br>3 (6834-7268) | Replicase<br>CP<br>MP<br>NABP | 2084<br>167<br>373<br>144 | DiVB-polyprot/0.0/93<br>ObRV1-MP/9e-66/80<br>CMLV-P4/3e-04/46 |
| *Betaflexiviridae/*<br>*Divavirus* | Korean chestnut<br>(*Castanea*<br>*crenata*); dicot | Korean<br>chestnut<br>virus 1 | KoChV1 | PRJNA534190 | Park, E.J.,<br>South<br>Korea;<br>unpublished | 7348 | MW328731 | 1 (45-7040)<br>2 (5285-6484) | Replicase<br>CP<br>MP | 2160<br>171<br>399 | DiVA-polyprot/0.0/99<br>ObRV1-MP/3e-47/73 |
| *Betaflexiviridae/*<br>*Divavirus* | Asian cow-wheat<br>(*Melampyrum*<br>*roseum*); dicot | *Melampyru*<br>*m roseum*<br>virus 2 | MelRoV2 | PRJDB5395 | [92] | 7094 | MW328733 | 1 (50-6403)<br>2 (4723-5826)<br>3 (6426-6911) | Replicase<br>CP<br>MP<br>NABP | 1954<br>163<br>367<br>161 | DiVB-polyprot/0.0/93<br>DiVA-MP/3e-63/84<br>CMLV-P4/6e-04/41 |
| *Betaflexiviridae/*<br>*Divavirus* | Capulincillo oak<br>(*Quercus*<br>*castanea*); dicot | *Quercus*<br>*castanea*<br>virus 1 | QueCaV1 | PRJNA532454 | [93] | 7761 | MW328737 | 1 (30-7025)<br>2 (5282-6469) | Replicase<br>CP<br>MP | 2150<br>181<br>395 | DiVA-polyprot/0.0/100<br>DiVA-MP/1e-42/67 |
| *Betaflexiviridae/*<br>*Fivirus* | Avocado (*Persea*<br>*americana*); dicot | Avocado<br>virus 1 | AvoV1 | PRJNA391003 | [80] | 8002 | MW328739 | 1 (115-5499)<br>2 (5560-6498)<br>3 (6281-6922)<br>4 (6997-7815)<br>5 (7640-7978) | Replicase<br>MP<br>CP<br>P4<br>NABP | 1794<br>312<br>213<br>272<br>112 | GKSV-Rep/0.0/90<br>FLV1/2e-32/88<br>FLV1/4e-4/88<br>no hits<br>GKSV-NABP/0.009/84 |
| *Deltaflexiviridae/*<br>*Deltaflexivirus* | Blue agave<br>(*Agave*<br>*tequilana*);<br>monocot | *Agave*<br>*tequilana*<br>virus 3 | ATV3 | PRJNA193469 | [79] | 8565 | MW328745 | 1 (13-6405)<br>2 (6732-7043)<br>3 (7126-7425)<br>4 (7427-7951)<br>5 (7945-8487) | Replicase<br>P2<br>P3<br>P4<br>P5 | 2130<br>103<br>99<br>174<br>180 | FgDFV1-Rep/0.0/97<br>no hits<br>no hits<br>no hits<br>no hits |
| *Deltaflexiviridae/*<br>*Deltaflexivirus* | Sesame (*Sesamum*<br>*indicum*); dicot | Sesame<br>virus 1 | SesV1 | PRNJA74261 | [94] | 8362 | MW328747 | 1 (54-6350)<br>2 (6535-6732)<br>3 (6803-7099)<br>4 (7276-7788) | Replicase<br>P2<br>P3<br>P4 | 2098<br>65<br>98<br>170 | FgDFV1-Rep/0.0/99<br>no hits<br>no hits<br>FgDFV1-P4/2e-28/100 |
|  |  |  |  |  |  |  |  | 5 (7772-8248) | P5 | 158 | FgDFV1-P5/1e-07/97 |
| *Tymoviridae/*<br>*Marafivirus* | Dove-tree<br>(*Davidia*<br>*involucrata*);<br>dicot | *Davidia*<br>*involucrata*<br>virus 1 | DaInvV1 | PRJNA513477 | [95] | 7109 | MW328750 | 1 (164-6955) | Replicase<br>CP | 2073<br>190 | MsMV1-polyprot/0.0/93 |
| *Tymoviridae/*<br>*Marafivirus* | Dove-tree<br>(*Davidia*<br>*involucrata*);<br>dicot | *Davidia*<br>*involucrata*<br>virus 2 | DaInvV2 | PRJNA513477 | [95] | 6775 | MW328752 | 1 (126-6632) | Replicase<br>CP | 1971<br>197 | CSDaV-polyprot/0.0/97 |
| *Tymoviridae/*<br>*Maculavirus* | Watercress<br>(*Nasturtium*<br>*officinale*); dicot | Watercress<br>associated<br>virus 1 | WaCraV1 | PRJNA284126 | [96] | 6744 | MW328754 | 1 (253-5637)<br>2 (5505-6317)<br>3 (6229-6648) | Replicase<br>CP<br>P3 | 1794<br>270<br>139 | BmMLV-Rep/0.0/99<br>BmMLV-CP/5e-49/81<br>BmMLV-P3 /6e-13/64 |
| *Tymoviridae/*<br>*unclassified* | Kava (*Piper*<br>*methysticum*),<br>dicot | Kava virus 1 | KaV1 | PRJNA494686 | [97] | 9491 | MW328755 | 1 (218-7048)<br>2 (7121-8215)<br>3 (8016-9395) | Replicase<br>P2<br>P3 | 2276<br>364<br>459 | UTyV1-Rep/3e-62/46<br>no hits<br>no hits |
| *Tymoviridae/*<br>*unclassified* | Polish wheat<br>(*Triticum*<br>*polonicum*);<br>monocot | Polish wheat<br>virus 1 | PolWhV1 | PRJNA304266 | [98] | 7702 | MW328751 | 1 (99-6752)<br>2 (6756-7148)<br>3 (7093-7644) | Replicase<br>P2<br>P3 | 2217<br>130<br>183 | FgMTV1-Rep/0.0/75<br>no hits<br>no hits |
| Unassigned<br>/*Gratylivirus* | Yellow poplar<br>(*Liriodendron*<br>*tulipifera*); dicot | yellow<br>poplar virus<br>1 | YePoV1 | PRJNA556244 | [99] | 6069 | MW328760 | 1 (35-5443)<br>2 (5515-6057) | Replicase<br>CP | 1802<br>180 | PYSaV-Rep/0.0/94<br>PRSaV-CP/2e-50/98 |

\*partial sequence

### Sequence assembly and identification

The nucleotide (nt) raw sequence reads from each analyzed SRA experiment linked to the TSA projects returning tymovirids-like hits were downloaded and pre-processed by trimming and filtering with the Trimmomatic tool as implemented in http://www.usadellab.org/cms/?page=trimmomatic, and the resulting reads were assembled *de novo* with Trinity release v2.11.00 using standard parameters. The transcripts obtained from *de novo* transcriptome assembly were subjected to bulk local BLASTX searches (E-value < 1e^−5^) against the TSA database obtained hits and a Refseq virus protein database available at ftp://ftp.ncbi.nlm.nih.gov/refseq/release/viral/viral.1.protein.faa.gz. The resulting viral sequence hits of each bioproject were visually explored. TSA and de novo assembled overlapping transcripts corresponding to tymovirids were re assembled and curated by mapping of SRA reads to the obtained sequences using Bowtie2 available at http://bowtie-bio.sourceforge.net/bowtie2/index.shtml, which was used to calculate total and mean coverage and reads per million (RPM) of each assembled virus sequence.

### Bioinformatics tools and analyses

#### Sequence analyses

ORFs were predicted with ORFfinder (https://www.ncbi.nlm.nih.gov/orffinder/), functional domains and architecture of translated gene products were determined using InterPro (https://www.ebi.ac.uk/interpro/search/sequence-search) and MOTIF search (https://www.genome.jp/tools/motif/). Importin-α dependent nuclear localization signals were predicted using cNLS Mapper available at http://nls-mapper.iab.keio.ac.jp/, and transmembrane domains were predicted using the TMHMM version 2.0 tool (http://www.cbs.dtu.dk/services/TMHMM/).

#### Percent identity matrices

Percentage amino acid (aa) sequence identity of the predicted ORFs of each virus identified in this study based on available tymovirids genome sequences were calculated using https://www.ebi.ac.uk/Tools/psa/emboss_needle/.

#### Phylogenetic analysis

Phylogenetic analysis based on the predicted replicase and capsid protein of the tymovirids detailed in Table S1, was done using MAFFT 7 https://mafft.cbrc.jp/alignment/software with multiple aa sequence alignments using FFT-NS-i as the best-fit model. The aligned aa sequences were used as input in MegaX software [25] to generate phylogenetic trees by the maximum-likelihood method using the best fit method which is detailed in the legend describing each phylogenetic tree. Local support values were computed using bootstrap with 1,000 replicates.

#### Recombination analysis

The sequences assembled in this study were examined for potential recombination events using the recombination detection program (RDP) version 5.05 with default settings [26]. Seven algorithms, BootScan, RDP, SiScan, GENECONV, MaxChi, Chimaera and Philpro were employed with a bonferroni corrected *P* value cut-off of <0.05. An event was considered as positive if was identified by at least three or more algorithms.

## Results

### Summary of discovered tymovirid sequences

The complete coding regions of 29 novel tymovirids were identified, in addition, partial genomic sequences for two novel viruses were assembled. These viruses were associated with 27 plant host species (Table 1). Bioinformatic and source data of each of the 29 viral sequences, as well as the GenBank accession numbers and proposed classification are listed in Table 1; the summary of the assembly statistics of each virus of the tymovirid sequences identified from the transcriptome data available in the NCBI database are presented in Table S2. Based on phylogenetic relatedness, genome organization and sequence identity, the novel viruses were tentatively assigned to the established tymovirales families: *Alphaflexiviridae, Betaflexiviridae, Deltaflexiviridae*, and *Tymoviridae*. 10 herbaceous dicots, 8 herbaceous monocots, and 9 woody dicots plant were identified as putative hosts of the viruses described in this study. Most of the woody dicot plants (6/9) appeared to be infected by betaflexiviruses belonging to the *Trivirinae* subfamily (Table 1). The conserved domains of deduced proteins encoded by each tymovirid sequence were determined by predictive algorithms, and are shown in Table S3. Genomic architecture and evolutionary placement of the 29 discovered viruses are described below, grouped by affinity to members of the diverse families within the order *Tymovirales*.

#### Alphaflexiviridae

The complete coding region of six putative alphaflexiviruses, tentatively named *Agave tequilana* virus 2 (ATV2), *Cymodocea nodosa* virus 1 (CyNoV1), indian gooseberry virus 1 (InGoV1), ferula virus 1 (FerV1), peltate yam virus 1 (PelYV1) and *Yucca gloriosa* virus 1 (YuGlV1), were assembled in this study (Table 1). ATV2, InGoV1, FerV1, PelYV1 and YuGlV1 genomes have five ORFs in the order 5’Rep-TGB1-TGB2-TGB3-CP-3’ (Fig.1A); while CyNoV1 genome has two ORFs in the order 5’-Rep-MP-3’ (Fig. 1A), which is a distinctive genomic organization among plant alphaflexiviruses.

**Figure 1.**
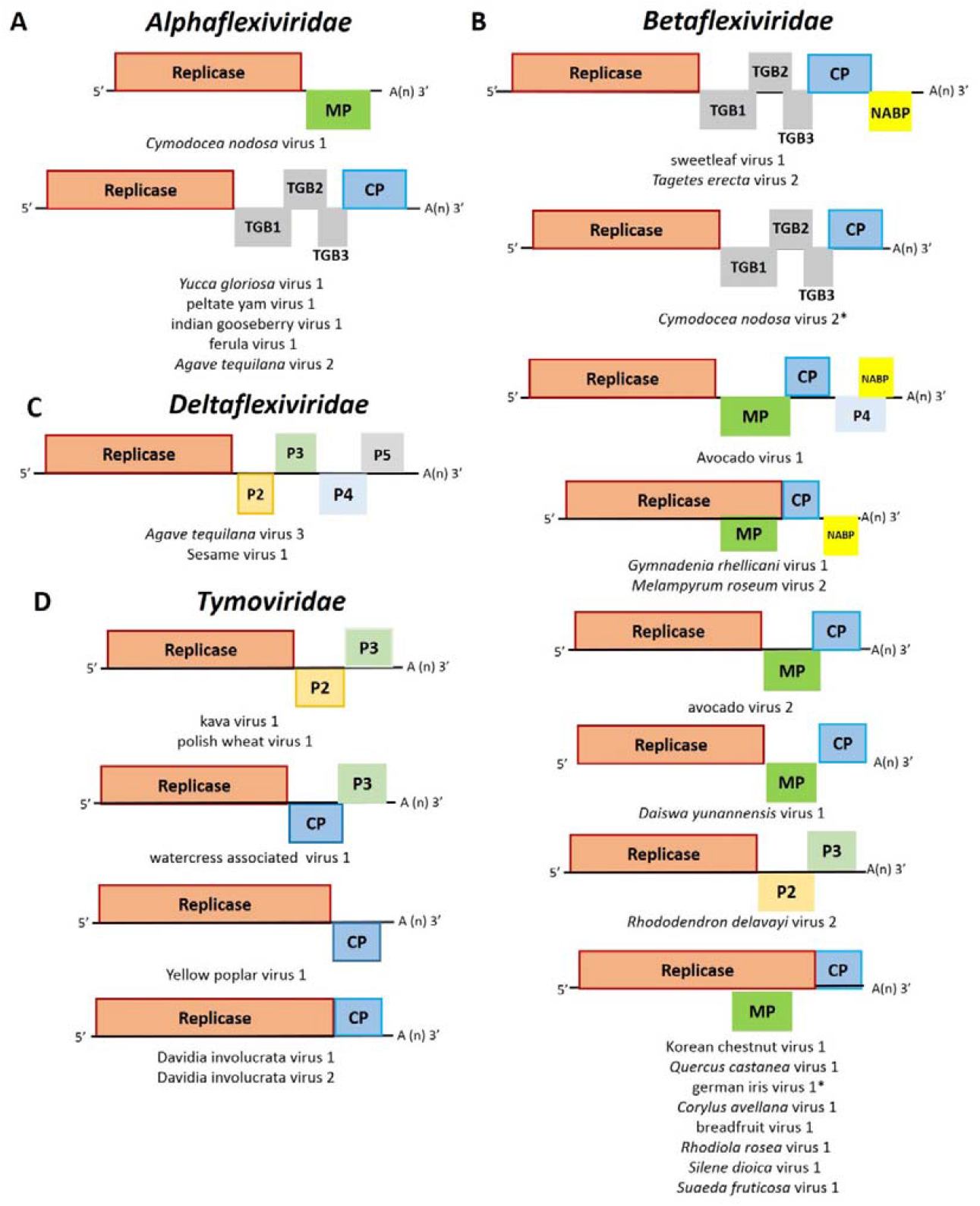
Diagrams depicting the genomic organization of each (A) alphaflexivirus (B) betaflexivirus, (C) deltaflexivirus and (D) tymovirus sequence identified in this study. Abbreviations: MP, movement protein CDS; TGB1, triple gene block protein 1 CDS; TGB2, triple gene block protein 2 CDS; TGB3, triple gene block protein 3 CDS; P capsid protein CDS; NABP, nucleic acid binding protein CDS.

Blast P searches showed that ATV2, InGoV1, FerV1, PelYV1 and YuGlV1 ORFs 1, 2, 3, 4 and 5 encoded proteins are related with replicases, TGB1, TGB2, TGB3 and capsid proteins encoded by other potexviruses (Table 1); while the BlastP search of CyNoV1 ORF1 and ORF2 encoded proteins showed that are related with the replicase and the MP encoded by the alphaflexivirus donkey orchid symptomless virus (DOSV) (Table 1).

The viral methyltransferase, viral helicase and the RdRP motifs were identified in the replicases encoded by ATV2, CyNoV1, InGoV1, FerV1, PelYV1 and YuGlV1. The 3A/RNA2 MP family conserved domain was identified in the MP encoded by the CyNoV1. Whereas the conserved domains viral helicase, plant viral MP, triple gene block 3 and flexi_CP conserved domains were identified in the TGB1, TGB2, TGB3 and CP, respectively, encoded by ATV2, InGoV1, FerV1, PelYV1 and YuGlV1 (Table S3).

Based on the ATV2, InGoV1, FerV1, PelYV1 and YuGlV1 aa sequences of the replicase and CP proteins, these viruses shared the highest sequence similarity with Vanilla virus X (VVX) (54.3%) and Cassava virus X (CsVX) (46.6%), Euonymus yellow mottle associated virus (EuYMaV) (58.6%) and EuYMaV (52.5%), Cnidium virus X (CnVX) (66.9%) and VVX (57.7%), Yam virus X (YVX) (57.8%) and YVX (50.1%), Potato virus X (PVX) (53.8%) and EuYMaV (47.2%), respectively. While, the aa sequence of the CyNoV1 replicase shared the highest sequence similarity with DOSV (50.1%). No recombination event was detected in the ATV2, CyNoV1, InGoV1, FerV1, PelYV1 and YuGlV1 genomic sequences.

In a phylogenetic tree based on the replicase aa sequence, PelYAV1 clustered together with the potexvirus ambrosia asymptomatic virus 1, InGoV1 clustered together with the potexvirus EuYMaV and FerV1 clustered together with the potexvirus CnVX, while ATV2 and YuGlV1 clustered together in a sister group to the potexviruses (Fig.2). Whereas, CyNoV1 clustered together with the platypuvirus DOSV (Fig.2). In the phylogenetric tree based on the CP aa sequences the same clustering were observed for ATV2, InGoV1, FerV1, PelYV1 and YuGlV1 (Figure S1).

**Figure 2.**
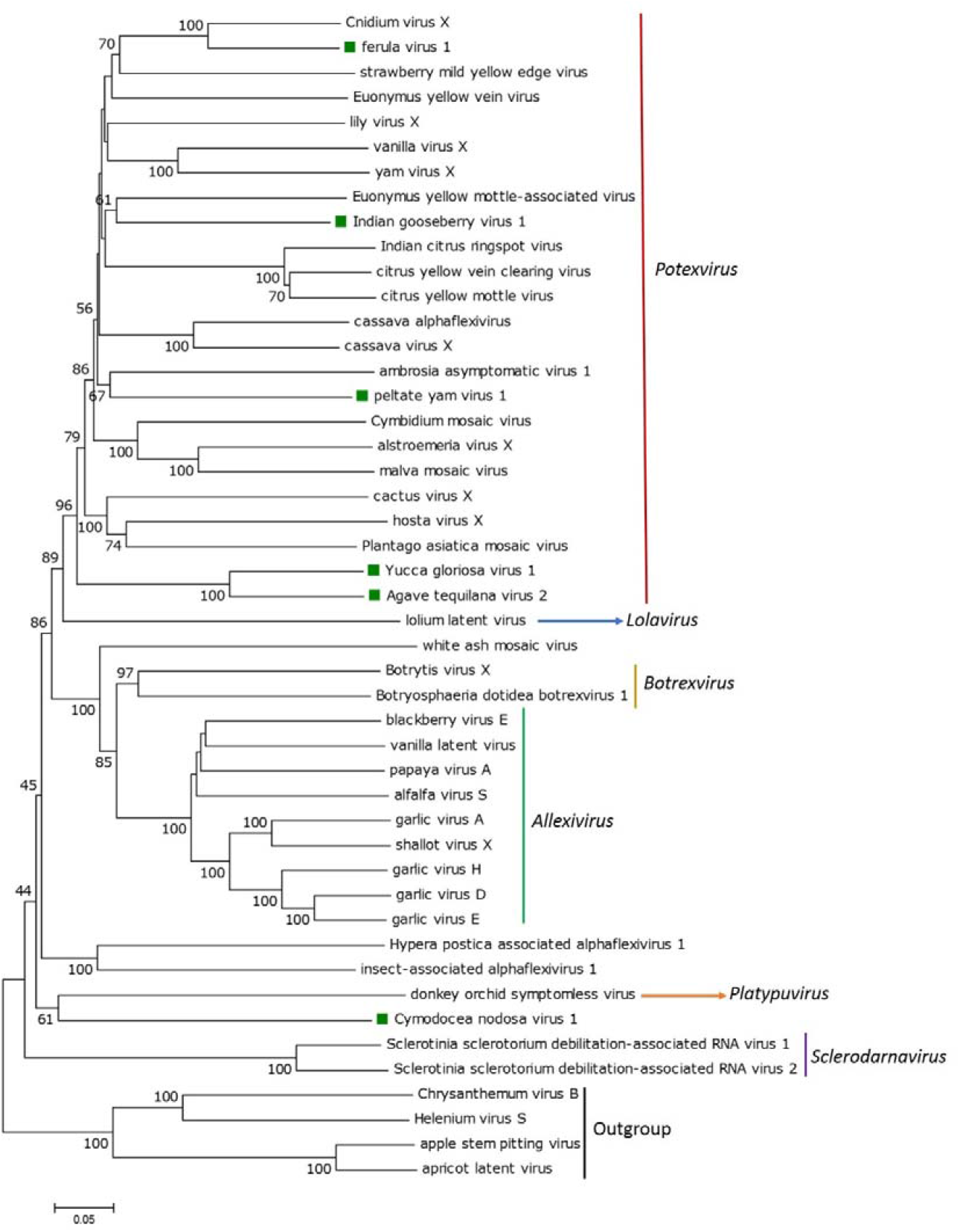
A Maximum Likelihood phylogenetic based on a multiple amino acid alignment of the replicase sequences of alphaflexiviruses, was constructed with the WAG+G+F model. Bootstrap values following 1000 replicates are given at the nodes, but only values above 40% are shown. Apple stem pitting virus, apricot latent virus, Helenium virus S and Chrysanthemum virus B replicase aa sequences were used as outgroup. The scale bar indicates the number of substitutions per site. Viruses identified in this study are noted with a green square. Accession numbers of every virus used to construct the ML tree are listed in Table S1.

#### Betaflexiviridae

The complete coding region of 15 putative betaflexiviruses, tentatively named as avocado virus 1 (AvoV1), avocado virus 2 (AvoV2), breadfruit virus 1 (BreFV1), *Corylus avellana* virus 1 (CorAvV1), *Daiswa yunannensis* virus 1 (DaYuV1), *Gymnadenia rhellicani* (GymRhV1), korean chestnut virus 1 (KoChV1), *Melampyrum roseum* virus 2 (MelRoV2), *Quercus castanea* virus 1 (QueCaV1), *Rhodiola rosea* virus 1 (RhRoV1), *Rhododendron delavayi* virus 2 (RhoDeV2), *Silene dioica* virus 1 (SiDiV1), *Suaeda fruticose* virus 1 (SuFruV1), sweetleaf virus 1 (SLeV1) and *Tagetes erecta* virus 2 (TaEV2), well as the partial genome of two other betaflexiviruses, named *Cymodocea nodosa* virus (CyNoV2) and German iris virus 2 (GerIV1), were assembled in this study (Table 1). BreFV1, CorAvV1, GerIV1, KoChV1, QueCaV1, RhRoV1, SiDiV1 and SuFruV1 genomes have two ORFs, named as Polyprotein-MP, where the ORF encoding the MP is overlapped (Fig.1B). GymRhV1 and MelRoV2 genomes have three ORFs in the order 5’-Polyprotein-MP-NABP-3’ where the ORF encoding the MP is overlapped (Fig.1B). AvoV2 and DaYuV1 genomes have three ORFs in the order 5’-Rep-MP-CP-3’ (Fig.1B), while RhoDeV2 has three ORFs in the order 5’-Rep-P2-P3-3’ (Fig.1B). AvoV1 genome has five ORFs in the order 5’-Rep-MP-CP-P4-NABP-3’ (Fig.1B). CyNoV2 genome has five ORFs in the order 5’-Rep-TGB1-TGB2-TGB3-CP-3’ (Fig.1B); while SleV1 and TaEV2 genome have six ORFs in the order 5’-Rep-TGB1-TGB2-TGB3-CP-NABP-3’ (Fig. 1B).

Blast P searches showed that BreFV1, CorAvV1, GerIV1, GymRhV1, KoChV1, MelRoV2, QueCaV1, RhRoV1, SiDiV1 and SuFruV1 ORFs 1 and 2 encoded proteins are similar to the polyprotein and movement proteins of other divaviruses, capilloviruses and unclassified betaflexiviruses (Table 1), while the GymRhV1 and MelRoV2 ORF3 showed identity with the NABPs of trichoviruses (Table 1). Blast P searches showed that AvoV2 and DaYuV1 ORFs 1, 2 and 3 encoded proteins are similar to the replicases, movement proteins and capsid proteins of other betaflexiviruses (Table 1); while Blast P searches showed that RhoDeV2 ORF1 encoded protein is more similar to the replicase of other trichoviruses, but no hits were found when ORFs 2 and 3 encoded proteins were searched (Table 1). Blast P searches showed that AvoV1 ORFs 1, 2, 3 and 5 encoded proteins are similar to the replicases, movement proteins, capsid proteins and nucleic acid binding proteins of other betaflexiviruses, whereas no hits were found when AvoV1 ORF4 encoded protein was subjected to a BlastP search (Table 1). Blast P searches showed that CyNoV2 ORFs 1, 2, 3, 4 and 5 encoded proteins are similar to the replicases, TGB1, TGB2, TGB3 and capsid proteins of other foveaviruses (for CyNoV2) (Table 1). BlastP searches showed that SleV1 and TaEV2 ORFs 1, 2, 3, 4, 5 and 6 encoded proteins are similar to the replicases, TGB1, TGB2, TGB3, capsid proteins and nucleic acid binding proteins of other carlaviruses (Table 1).

The viral methyltransferase, viral helicase and the RdRP motifs were identified in the replicases encoded by Avo1, Avo2, RhoDeV2, SLeV1 and TaEV2, and in the polyproteins encoded by BreFV1, CorAvV1, DaYuV1, GymRhV1, KoChV1, MelRoV2, QueCaV1, RhRoV1, SiDiV1 and SuFruV1. The viral methyltransferase motif was not found in the CyNoV2 encoded replicase and GerIV1 encoded polyprotein, probably because they were only partially assembled and its N-terminal region is missing, but the viral helicase and RdRP motifs were identified. Furthermore, the carlavirus endopeptidase conserved domain was found in the replicases encoded by SLeV1 and TaEV2. Whereas the trichovirus coat protein motif was found in the polyprotein encoded by BreFV1, CorAvV1, GerIV1, GymRhV1, KoChV1, MelRoV2, QueCaV1, RhRoV1, SiDiV1 and SuFruV1 and in the CP encoded by AvoV1, AvoV2 and DaYuV1; while the DUF1717 conserved domain was identified in the polyprotein encoded by BreFV1, CorAvV1, GerIV1, RhRoV1 and SiDiV1 and the 2OG-FeII_Oxy_2 motif was identified in the replicase encoded by AvoV2 and RhoDeV2. Moreover, the conserved domain viral MP was identified in the MP encoded by AvoV1, AvoV2, BreFV1, CorAvV1, DaYuV1 GerIV1, GymRhV1, KoChV1, MelRoV2, QueCaV1, RhRoV1, SiDiV1 and SuFruV1. Whereas the conserved domains viral helicase, plant viral MP, triple gene block 3 and flexi_CP conserved domains were identified in the TGB1, TGB2, TGB3 and CP, respectively, encoded by CyNoV2, SleV1 and TaEV2. Furthermore, the Carlavirus coat motif was identified in the CP encoded by SLeV1 and TaEV2, and the carlavirus putative NABP conserved domain was identified in the NABP encoded by AvoV1, SLeV1 and TaEV2 (Table S3), while a viral_NABP conserved domain was identified in the NABP encoded by GymRhV1 and MelRoV2 (Table S3). No conserved domains were identified in the proteins 2 and 3 encoded by RhoDeV2, and in the protein 4 encoded by AvoV1 (Table S3), but a bipartite NLS was detected in the P3 encoded by RhoDeV2 and in the P4 encoded by AvoV1.

Based on the AvoV1, AvoV2, CyNoV2, DaYuV1, SleV1 and TaEV2 aa sequences of the replicase and CP proteins, these viruses shared the highest sequence similarity with grapevine Kizil Sapak virus (GKSV) (50.1%) and fig latent virus 1 (FLV1) (48.6%), citrus leaf blotch virus (CLBV) (55.6%), and Camellia ringspot-associated virus 2 (CRSaV2) (52.8%), Asian prunus virus 3 (APV3) (43.0%), and Grapevine rupestris stem pitting-associated virus (GRSPaV) (44.7%), Agapanthus virus A (AgVA) (54.4%) and AgVA (51.7%), Helenium virus S (HelVS) (60.6%) and HelVS (63.9%), HelVS (60.9%) and (HelVs) (65.7%), respectively. Whereas the aa sequence of the BreFV1, CorAvV1, GerIV1, GymRhV1, KoChV1, MelRoV2, QueCaV1, RhRoV1, RhoDeV2, SiDiV1 and SuFruV1 replicase protein, shared the highest sequence similarity with Apple stem grooving virus (ASGV) (69.4%), ASGV (58.8%), ASGV (53.6%), diuris virus B (DiVB) (52.2%), diuris virus A (DiVA) (53.5%), DiVB (47.7%), DiVB (52.7%), Hobart betaflexivirus 1 (HoBFV1) (47.1%), cherry mottle leaf virus (CMLV) (44.1%), ASGV (49.7%) and Camellia ringspot-associated virus 3 (CRSaV3) (52.8%), respectively. No recombination event was detected in the betaflexiviruses assembled in this study.

In a phylogenetic tree based on the replicase aa sequence of viruses belonging to the subfamily *Quinvirinae*, SleV1 and TaEV2 clustered together and this group was related with the carlavirus HelVS. On the other hand, CyNoV2 formed a monophyletic group that was grouped with the unclassified betaflexiviruses yam virus Y and sugarcane striate mosaic-associated virus (Fig. 3). Whereas, in the phylogenetic tree based on the CP aa sequence, SleV1 and TaEV2 clustered together and this group was related with the carlavirus HelVS; while CyNoV2 clustered together with the foveavirus rubus virus 1 (Figure S2).

**Figure 3.**
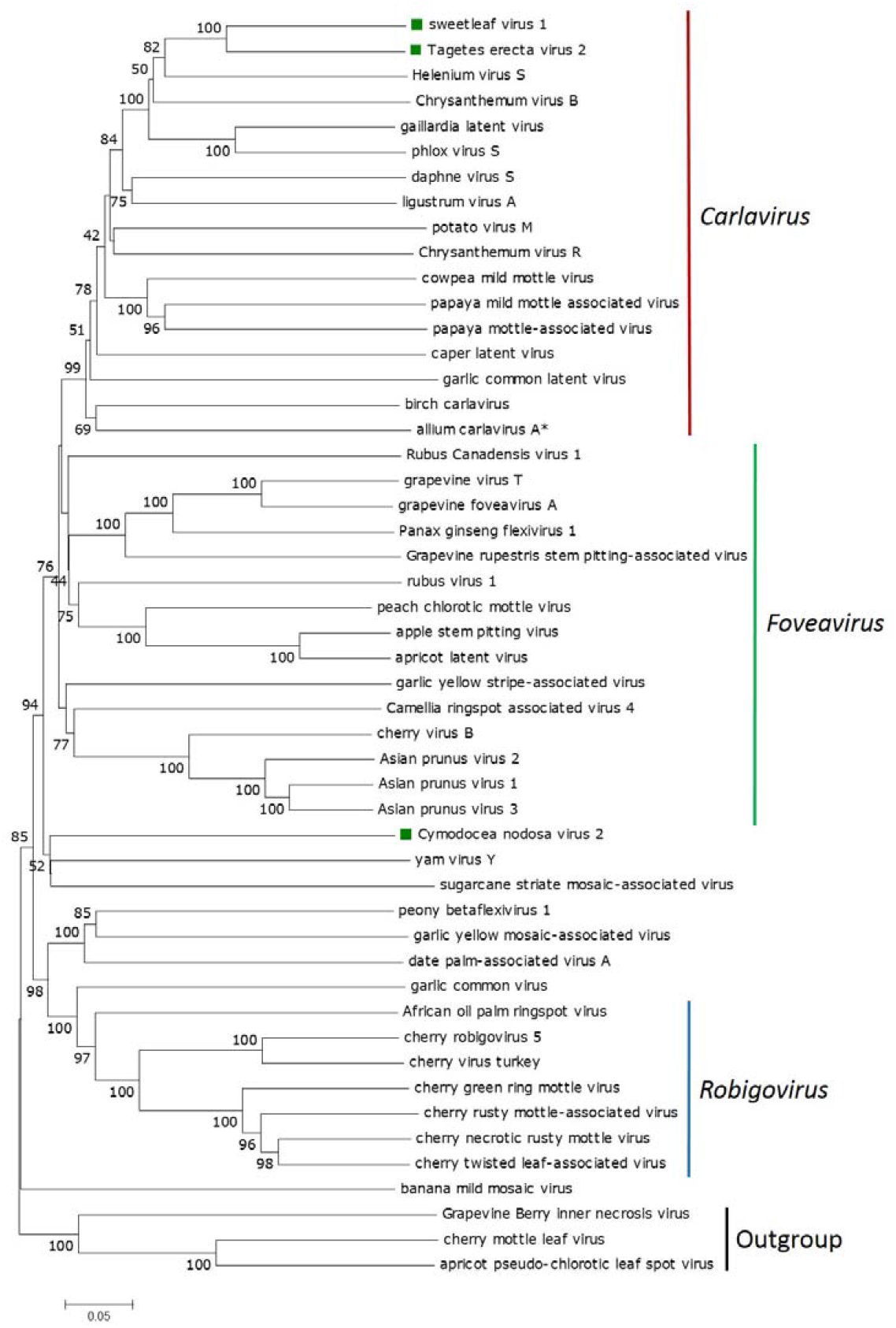
A Maximum Likelihood phylogenetic based on a multiple amino acid alignment of the replicases sequences of viruses of the subfamily *Quinvirinae*, was constructed with the LG+G+F model. Bootstrap values following 1000 replicates are given at the nodes, but only values above 40% are shown. Apricot pseudo-chlorotic leaf spot virus, cherry mottle leaf virus and Grapevine Berry Inner necrosis virus replicases aa sequences were used as outgroup. The scale bar indicates the number of substitutions per site. Viruses identified in this study are noted with a green square. Viruses identified with an * correspond to unpublished sequences derived from the analysis of RNA-Seq data publically available at the NCBI. Accession numbers of every virus used to construct the ML tree are listed in Table S1.

In a phylogenetic tree based on the replicase aa sequence of viruses belonging to the subfamily *Trinvirinae*, AvoV1 was placed is the same clade as GKSV and FLV1, that is separated from the clades formed by other genera belonging to the *Trinvirinae* subfamily (Fig.4). RhoDeV2 formed a monophyletic cluster that is related with the *Chordovirus* clade and the ravavirus Ribes americanum virus A (Fig. 4).

**Figure 4.**
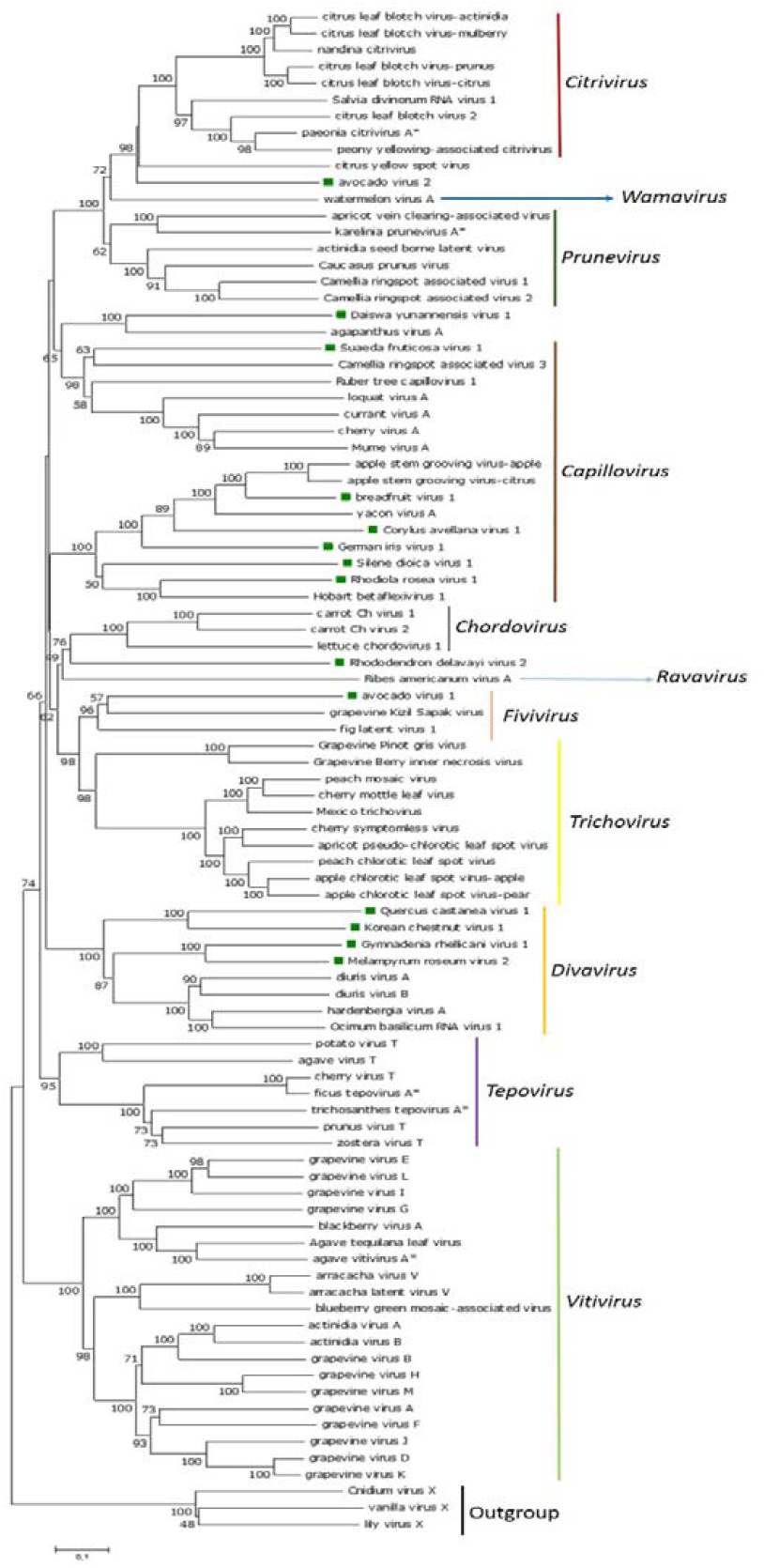
A Maximum Likelihood phylogenetic based on a multiple amino acid alignment of the replicases sequences of viruses of the subfamily *Trinvirinae*, was constructed with the LG+G+F model. Bootstrap values following 1000 replicates are given at the nodes, but only values above 40% are shown. Cnidium virus X, lily virus X and vanilla virus X replicases aa sequences were used as outgroup. The scale bar indicates the number of substitutions per site. Viruses identified in this study are noted with a green square. Viruses identified with an * correspond to unpublished sequences derived from the analysis of RNA-Seq data publically available at the NCBI. Accession numbers of every virus used to construct the ML tree are listed in Table S1.

GymRhV1 clustered together with MelRoV2, and KoChV1 with QueCaV1 and this two clades were grouped with the *Divavirus* clade (Fig. 4). AvoV2 formed a monophyletic cluster that is related with the *Citrivirus* clade and the unclassified betaflexivirus citrus yellow spot virus (Fig. 4). DaYuV1 clustered together with AgVA and this cluster is separated from the clades formed by members of other genera belonging to the *Trinvirinae* subfamily (Fig. 4). SuFruV1 clustered together with CRSaV3 and this clade grouped with the capilloviruses (Fig, 4). RhRoV1 clustered together with HoBFV1 and SiDiV1 and this cluster grouped with capilloviruses (Fig. 4). Whereas, BreFV1, CorAvV1 and GerIV1 clustered with the capilloviruses ASGV and yacon virus A (Fig. 4). In the phylogenetic tree based on the CP aa sequence, AvoV1 was placed in the same clade as GKSV and FLV1 (Figure S3). GymRhV1 clustered together with MelRoV2, and KoChV1 with QueCaV1 and this two clades were grouped with the *Divavirus* clade (Figure S3). AvoV2 formed a monophyletic cluster that is related with the *Prunevirus* clade (Supp. Figure 3). DaYuV1 clustered together with AgVA (Figure S3). SuFruV1 is related with CRSaV3 and capilloviruses (Figure S3). RhRoV1 clustered together with HoBFV1 and SiDiV1 and this cluster grouped with known capilloviruses, while, CorAvV1 and GerIV1 clustered with the capilloviruses yacon virus A and BreFV1 with the capillovirus ASGV (Figure S3).

#### Deltaflexiviridae

The complete coding region of two putative deltaflexiviruses, tentatively named Agave tequilana virus 3 (ATV3) and sesame virus 1 (SesV1), were assembled in this study (Table 1). ATV3 and SesV1 genomes consist of five ORFs in the order 5’Rep-P2-P3-P4-P5-3’ (Fig.1C).

Blast P searches showed that ATV3 and SesV1 ORF1 encoded proteins are similar to the replicases of other deltaflexiviruses (Table 1); while BlastP searches showed SesV1 ORFs 4 and 5 are similar to the P4 and P5 proteins of other deltaflexiviruses (Table 1). No hits were found when ATV3 ORFs 2, 3, 4 and 5 encoded proteins, as well as SesV1 ORFs 2, 3 encoded proteins, were subjected to a BlastP search (Table 1).

The typical viral methyltransferase and the RdRP motifs were identified in the replicase encoded by ATV3, and these two conserved domains as well a viral helicase conserved domain was identified in the SesV1 encoded replicase; while no conserved domains were identified in the other proteins encoded by ATV3 and SesV1 (Table S3).

Based on the ATV3 and SesV1 aa sequences of the replicase, these viruses shared the highest sequence similarity with Fusarium graminearum deltaflexivirus 1 (FgDFV1) (55.7%) and FgDFV1 (58.2%), respectively.

No recombination event was detected in the deltaflexiviruses assembled in this study.

In a phylogenetic tree based on the replicase aa sequence, ATV3 and SesV1 were grouped with other known deltaflexiviruses (Fig.5).

**Figure 5.**
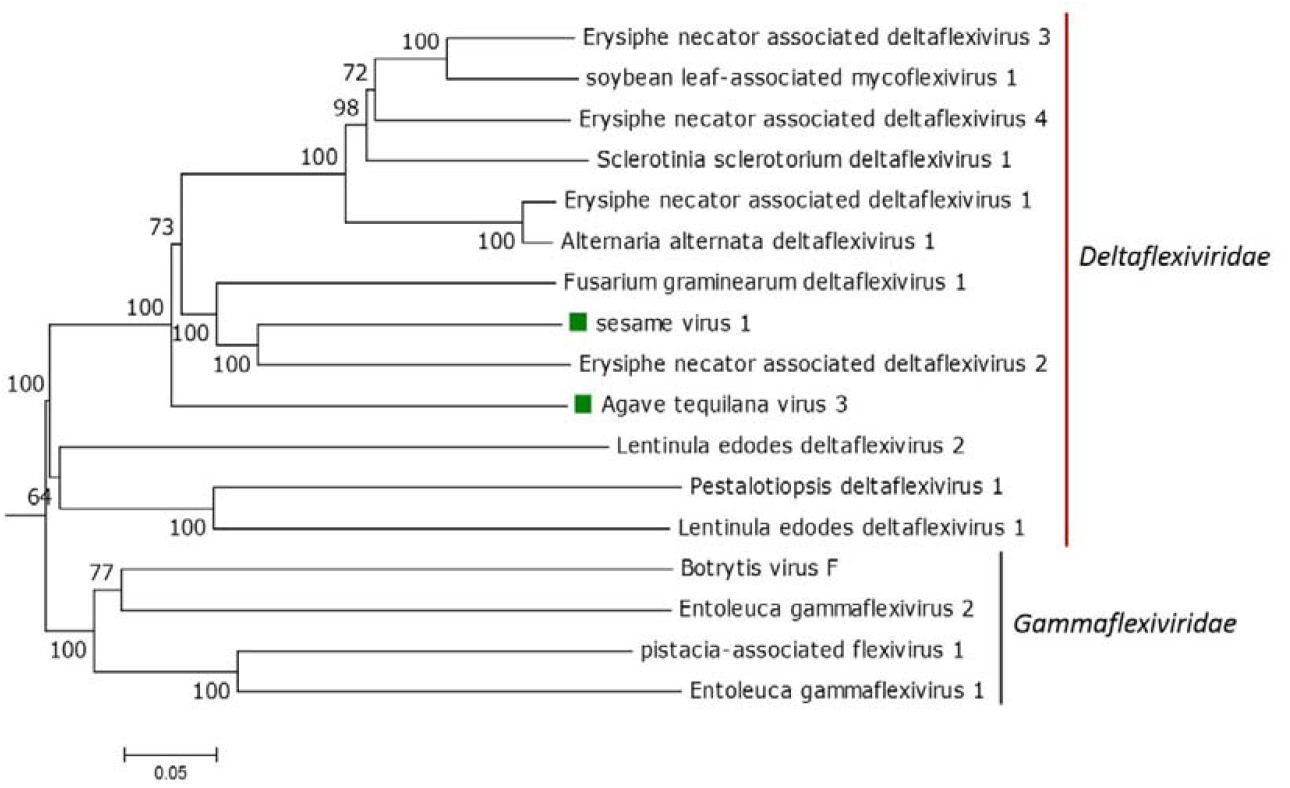
A Maximum Likelihood phylogenetic based on a multiple amino acid alignment of the replicase sequences of deltaflexiviruses, was constructed with the JTT+G model. Bootstrap values following 1000 replicates are given at the nodes, but only values above 40% are shown. The gammaflexiviruses Botrytis virus F, Entoleuca gammaflexivirus 1, Entoleuca gammaflexivirus 2 and pistacia-associated flexivirus 1 replicases aa sequences were used as outgroup. The scale bar indicates the number of substitutions per site. Viruses identified in this study are noted with a green square. Accession numbers of every virus used to construct the ML tree are listed in Table S1.

#### Tymoviridae and tymo-like virus

The complete coding region of six putative tymoviruses, tentatively named *Davidia involucrata* virus 1 (DaInvV1), *Davidia involucrata* virus 2 (DaInvV2), kava virus 1 (KaV1), polish wheat virus 1 (PolWh1), watercress associated virus 1 (WaCraV1) and yellow poplar virus 1 (YePoV1) were assembled in this study (Table 1). DaInv1 and DaInv2 genomes have one ORF (Fig.1D), YePoV1 has two ORFs in the order 5’Rep-CP-3’ (Fig.1D); while KaV1 and PolWhV1 genomes have three ORFs in the order 5’-Rep-P2-P3-3’, while WaCraV1 genome has three ORFs in the order 5’-Rep-CP-P3-3’ (Fig. 1D).

Blast P searches showed that DaInv1 and DaInv2 ORF encoded protein are similar to the polyprotein of other marafiviruses (Table 1). Blast P searches showed that KaV1, PolWhV1, WaCraV1 and YePoV1ORF1 encoded proteins are similar to the replicases of other tymo-like viruses, while WaCraV1 and YePoV1 ORF 2 encoded proteins are similar to the CP of other tymo-like viruses, and WaCraV1 ORF3 encoded protein is similar to the P3 encoded by other tymo-like viruses (Table 1). No hits were found when KaV1, PolWhV1 and WaCraV1 ORFs 2 and 3 encoded proteins were subjected to a BlastP search (Table 1).

The viral methyltransferase, viral helicase and the RdRP motifs were identified in the DaInV1 and DaInv2 polyproteins and in the replicase encoded by KaV1, PolWhV1, WaCraV1 and YePoV1, while the tymovirus endopeptidase conserved domain was identified in the DaInV1 and DaInv2 polyproteins and in the replicase encoded by PolWhV1 and WaCraV1. Whereas the tymovirus coat protein conserved domain was identified in the DaInV1 and DaInv2 polyproteins and in the capsid protein encoded by WaCraV1 (Table S3). Moreover, a transmembrane domain was identified in the protein 2 and protein 3 encoded by KaV1 and WaCraV1, respectively.

Based on the nearly complete assembled genomes of DaInv1, DaInv2, KaV1, PolWhV1, WaCraV1 and YePoV1, these viruses shared the highest sequence similarity with Medicago sativa marafivirus 1 (MsMV1) (57.2%), citrus sudden death-associated virus (CSDaV) (59.1%), Ullucus tymovirus 1 (UTyV1) (42.3%), Fusarium graminearum mycotymovirus 1 (FgMTV1) (51.6%), bee macula-like virus 2 (BeeMLV2) (55.2%) and prunus yellow spot-associated virus (PYSaV) (59.7%), respectively. While the DaInv1, DaInv2, WaCraV1 and YePoV1 CP shared the highest amino acid sequence identity with MsMV1 (57.36%), CSDaV (59.09%), BeeMLV2 (48.70%) and PYSaV (44.94%) No recombination events were detected in the DaInV1, DaInV2, KaV1, PolWhV1, WaCraV1 and YePoV1 genomic sequences.

In a phylogenetic tree based on the replicase aa sequence, DaINV1 and DaINV2 clustered with marafiviruses. DaINV1 was related to two clades, one formed by Medicago sativa marafivirus 1 and alfalfa virus F, and the other by citrus virus C and grapevine red globe virus; while DaInv2 was closely related with blackberry virus S (Fig.6). WaCraV1 clustered together with the maculaviruses BeeMLV2 and Bombyx mori macula-like virus. KaV1 formed a monophyletic cluster that was distantly related with known tymoviruses, while PolWhV1 clustered together with the mycotymoviruses FgMTV1 and this viruses grouped with the cluster formed by the mycotymoviruses Sclerotinia sclerotorium mycotymovirus 1, Sclerotinia sclerotorium mycotymovirus 2 and Botrytis cinerea mycotymovirus 1 (Fig.6). Whereas, YePoV1 clustered together with the tymo-like viruses PYSaV and grapevine associated tymo-like virus (GaTLV) (Fig.6).

**Figure 6.**
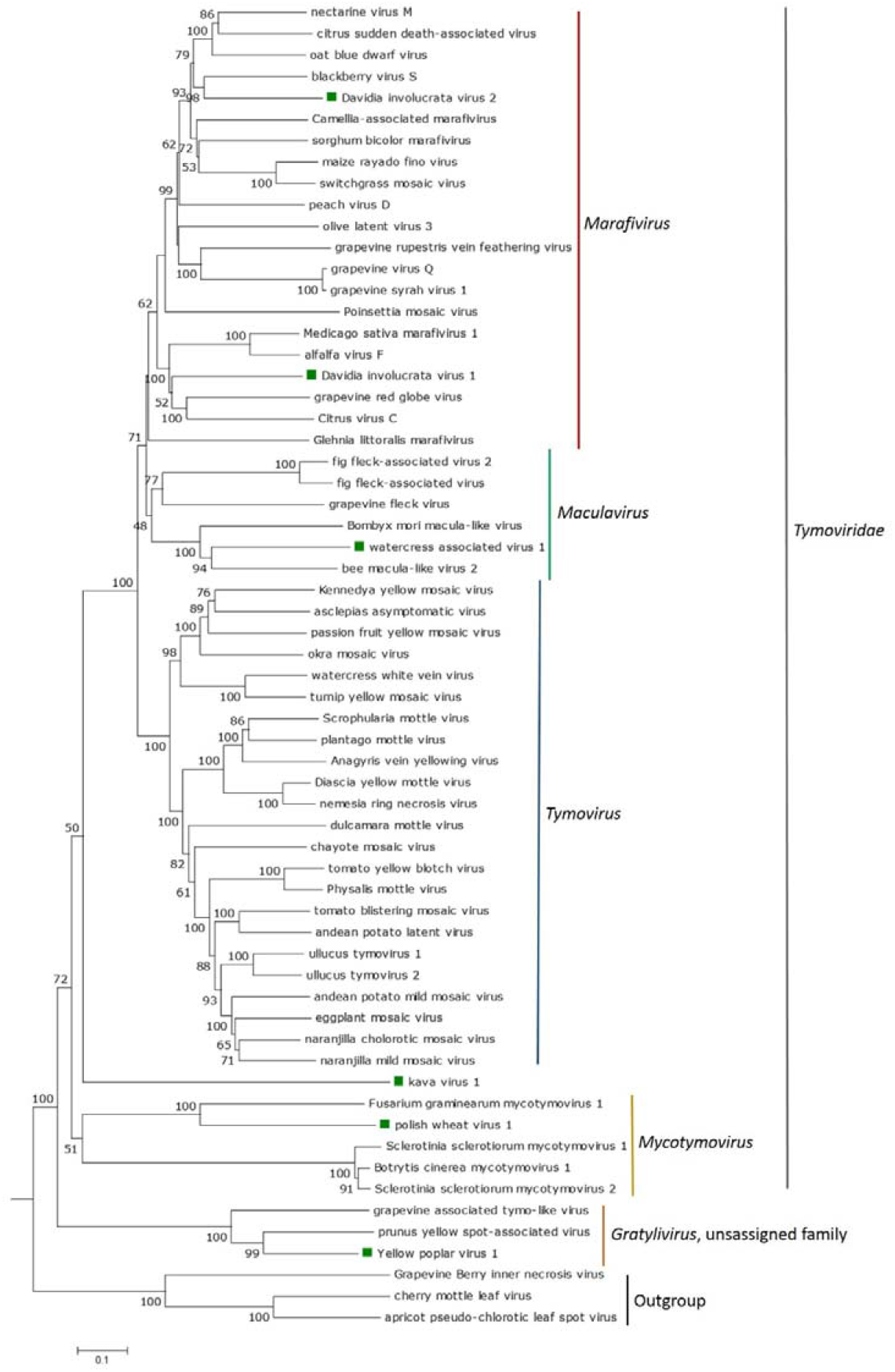
A Maximum Likelihood phylogenetic based on a multiple amino acid alignment of the replicase sequences of tymovirids, was constructed with the WAG+G+F model. Bootstrap values following 1000 replicates are given at the nodes, but only values above 40% are shown. . Apricot pseudo-chlorotic leaf spot virus, cherry mottle leaf virus and Grapevine Berry Inner necrosis virus replicases aa sequences were used as outgroup. The scale bar indicates the number of substitutions per site. Viruses identified in this study are noted with a green squares. Accession numbers of every virus used to construct the ML tree are listed in Table S1.

## Discussion

In the last few years, the employment of HTS, allowed the identification and characterization of several novel viruses associated to the order *Tymovirales*, which may be asymptomatic [27–33]. Moreover, some tymovirids were also discovered mining publically available transcriptome datasets at the NCBI [34–40]. Thus, many tymovirales-like sequences could be hidden in publicly available plant transcriptome datasets. Therefore, we conducted an extensive search of tymovirales-like sequences querying the transcriptome shotgun assembly (TSA) which resulted in the identification of 31 novel tymovirids. After extensive structural and functional annotation and evolutionary analyses we suggest these viruses correspond to potential new members of new virus species, which expand the diversity of the complete *Tymovirales* order by ca. 30%.

### Conserved domains

All the replicase proteins assembled in this study contained the methyltransferase (MTR), RNA helicase (Hel), and RNA-dependent RNA polymerase (RdRp) domains. These domains have been identified in a wide range of ssRNA viruses, where MTR is involved in capping, which enhances mRNA stability [41], the RdRp corresponds to the main viral replicase, an essential protein encoded in all RNA viruses, while the HEL domain plays multiple roles at different stages of viral RNA replication [42].

Transmembrane domains were identified in the TGB2 and TGB3 proteins encoded by ATV2, CyNoV2, FerV1, InGoV1, PelYV1, SLeV1, TaEV2 and YuGlV1, which also were reported for the TGB2 and TGB3 proteins encoded by other foveaviruses, carlaviruses, potexviruses, robigoviruses and allexiviruses [43], thus supporting the membrane-association and involvement in viral movement of these two proteins [44–46].

### Genome organization and phylogenetic relationships of the novel tymovirids

#### Alphaflexiviridae

ATV2, InGoV1, FerV1, PelYV1 and YuGlV1 showed a genomic organization 5’-Rep-TGB1-TGB2-TGB3-CP-3’, that is typical of potexviruses [16] and these five viruses are phylogenetically related with other potexviruses, but the highest nt identities when the Rep and CP genes were compared with other potexviruses was below 72%, which is the criteria to demarcate species in the *Alphaflexiviridae* family [16]. Thus, based on sequence similarities, genomic organization and phylogenetic relationships, ATV2, InGoV1, FerV1, PelYV1 and YuGlV1 should be classified as new members of the genus *Potexvirus* in the family *Alphaflexiviridae*.

On the other hand, CyNoV1 has a distinctive genomic organization among alphaflexiviruses because only two ORFs, encoding a replicase and MP, were identified in its genome, likely being the first alphaflexivirus reported so far that has only two ORFs. Like the sclerodarnaviruses, CyNoV1 does not encode a putative CP. Moreover, its replicase and MP are distantly related with that one encoded by the platypuvirus DOSV. Similarly to the DOSV MP, the CyNoV1 encoded MP is related with the 3A-like MPs of viruses belonging to the families *Tombusviridae* and *Virgaviridae* [47], which is distinctive among alphaflexiviruses, where the TGB proteins are the MPs [16]. Despite that the replicase nt sequence similarity between CyNoV1 and DOSV is above the 45% threshold value to demarcate genus in the family *Alphaflexiviridae* [16], and that the Rep of these two viruses grouped together in the phylogenetic tree, CyNoV1 has a genomic organization different than that one reported for DOSV [47]; thus CyNoV1 could be classified as a member of a novel genus belonging to the family *Alphaflexiviridae*.

#### Betaflexiviridae

SleV1 and TaEV2 showed a genomic organization 5’-Rep-TGB1-TGB2-TGB3-CP-NABP-3’, that is typical of carlaviruses [15]. Moreover, these two viruses are phylogenetically related with other carlaviruses, but the highest nt identities when the Rep and CP genes were compared with other carlaviruses was well below 72%, which is the criteria to demarcate species in the *Betaflexivirus* family [15]. Thus, based on sequence similarities, genomic organization and phylogenetic relationships, SleV1 and TaEV2 should be classified as novel members of the genus *Carlavirus* in the family *Betaflexiviridae*. BreFV1, CorAvV1, GerIV1, RhRoV1, SiDiV1 and SuFruV1 showed a genomic organization with two overlapping ORFs; where one encodes a Rep protein fused to the CP, while the other encodes a MP, which is typical of capilloviruses [30]. The highest nt identity when the Rep and CP proteins were compared with other betaflexiviruses was below 72%, which is the criteria to demarcate species in the *Betaflexiviridae* family [15]. The BreFV1, CorAvV1, GerIV1, RhRoV1, SiDiV1 Rep and CP clustered together with known capilloviruses, while SuFruV1 Rep and CP was similar to the one of CRSaV3 and these two viruses are related with known capilloviruses. Interestingly, the CRSaV3 genomic organization is different than the one of SuFruV1 and known capilloviruses [48]. Moreover, the conserved domain DUF1717 was identified in the polyprotein of BreFV1, CorAvV1, GerIV1, RhRoV1 and SiDiV1, but not in the SuFruV1 polyprotein. The DUF1717 domain, which function is unknown, was found in the polyprotein of the capilloviruses ASGV and cherry virus A [49, 50] as well as in the polyprotein of the unclassified betaflexivirus HoBFV1; however this domain was not found in the polyprotein encoded by CRSaV3. Sequence similarities, genomic organization and phylogenetic relationships supports that BreFV1, CorAvV1, GerIV1 RhRoV1, SiDiV1, and SuFruV1 as well as the previously reported HoBFV1 (Roberts et al., 2018) and CRSaV3 [48] should be classified as members of the genus *Capillovirus* in the family *Betaflexiviridae*, where CRSaV3 would be the first capillovirus with three ORFs instead of two, while HoBFV1 is likely the first insect-associated capillovirus described so far.

KoChV1 and QueCaV1 genomes have two overlapping ORFs, one is the fused Rep-CP ORF and the other one encodes the MP, which is typical of divaviruses [34]; while GymRhV1 and MelRoV2 also have a third ORF which encodes a putative NABP. These four viruses were distantly related with other divaviruses, and the highest nt identity when the Rep and CP genes were compared with other betaflexiviruses was below 72%, which is the criteria to demarcate species in the *Betaflexiviridae* family [15]. Moreover, these viruses clustered together with divaviruses both in the Rep and CP phylogenetic trees, thus they likely share a similar evolutionary history. Thus, based on sequence similarities, genomic organization and phylogenetic relationships, KoChV1 and QueCaV1 should be classified as novel members of the genus *Divavirus* in the family *Betaflexiviridae*. Despite that GymRhV1 and MelRoV2 have a different genomic organization than known divaviruses, based on sequence similarities and phylogenetic relationships these two viruses should also be classified as novel members of the genus *Divavirus*. The genus *Trichovirus* also contains viruses that display different genomic organizations [15], which support the potential classification of GymRhV1 and MelRoV2 as divaviruses.

The genomic organization of AvoV1 5’-Rep-MP-CP-P4-NABP-3’, is similar to that reported for the betaflexivirus GKSV [52]. The highest nt identity when the Rep and CP proteins were compared with other betaflexiviruses was below 72%, which is the criteria to demarcate species in the *Betaflexiviridae* family [15]. Neither conserved domains were identified, nor any BlastP hits were obtained when the AvoV1 ORF4 encoded protein was analyzed, which is similar to that reported for the encoded protein by the GKSV ORF4 [52]. Interestingly, in this protein of unknown function, a NLS was found in the one encoded by AvoV1, but no NLS was detected in the protein encoded by GKSV ORF4. Further studies should be conducted to characterize the putative function of this protein. AvoV1 and GKSV, along with FLV1, which full sequence has not been obtained yet [53], were phylogenetically related in a distinct clade from other genera belonging to the subfamily *Trivirinae*. GKSV and FLV1 were proposed to belong to a new genus named as *Fivivirus* [52]. Based on sequence similarities, genomic organization and phylogenetic relationships, AvoV1 should be classified as novel member of this genus, which name could be changed to Avofivivirus (derived from **Avo**cado-**Fi**cus-**Vi**tis-virus).

CyNoV2 genome organization 5’-Rep-TGB1-TGB2-TGB3-CP-3’, resembles the one described both for foveaviruses [54] and for the unclassified betaflexiviruses banana mild mosaic virus [55], sugarcane striate mosaic-associated virus [56] and yam virus Y [57]. The identities between the nt sequence of the replicase and CP of CyNoV2 with other betaflexiviruses is below 45%, that is the genus demarcation criteria for betaflexiviruses [15]. CyNoV2 replicase clustered together with the unassigned betaflexiviruses sugarcane striate mosaic-associated virus and yam virus X, while its CP was phylogenetically related with known foveaviruses. Based on sequence similarities, and the incongruences in the phylogenetic relationships observed between Rep and CP proteins, which likely indicates a distinct evolutionary history of CyNoV2 Rep and CP, this virus may represents a novel member of a new genus within the *Quinvirinae* subfamily in the *Betaflexiviridae* family, which could be tentatively named as Cynovirus.

The AvoV2 genome has three ORFs with a genomic organization 5’-Rep-MP-CP-3’, that is similar to that reported for the unclassified betaflexvirus citrus yellow spot virus [58], almost identical to that of citriviruses, where the ORF that encodes the MP is located adjacent to the ORF that encodes the Rep protein [15]. However, in the AvoV2 genomes, the ORF encoding the CP is overlapped with that one encoding the MP [58]; while in the citriviruses these two ORFs are separated by an intergenic region [15, 32, 59]. Similarly to the AvoV2 genome, in the pruneviruses the ORFs encoding the MP and CP are overlapped [15]; however most of the pruneviruses have four ORFs instead of three [48]. AvoV2 Rep and CP were distantly related with other citrivirus and pruneviruses, respectively, and the replicase and CP nt sequence similarity is above the 45% threshold value to demarcate genus in the family *Betaflexiviridae* [15]. However, this virus formed a monophyletic cluster that was related with citrus yellow spot and the *Citrivirus* clade, and the *Prunevirus* clade when the Rep and CP proteins were analyzed, respectively. Therefore, there is incongruences in the phylogenetic relationships observed between Rep and CP proteins, which likely indicates a distinct evolutionary history of AvoV2 Rep and CP. Thus, based on its genomic organization and phylogenetic relationships, AvoV2 should be classified as a novel member of a new genus within the subfamily *Trivirinae* in the family *Betaflexiviridae*, which could be tentatively named as Avovirus.

It is tempting to speculate that the discrepancy regarding phylogenetic tree topology between the one inferred from the Rep sequences and the one inferred from the CP sequences found for CyNoV2 and Avo2 could be linked to ancestral recombination events, or that these genes have different mutation rates. Similar hypotheses were speculated for other betaflexiviruses where discrepancies on the REP and CP evolutionary relationships have also been observed [58, 60].

The DaYuV1 genome has three ORFs, with a genomic organization 5’-Rep-MP-CP-3’, that is similar to the unclassified betaflexivirus AgVA, and reminiscent to the one reported for those trichoviruses with three ORFs, where the ORF that encodes the MP is slightly overlapped with the ORF that encodes the Rep [15]. However, in the DaYuV1 and AgVA genomes the ORF encoding the CP is separated from the ORF encoding the MP by a short intergenic region, while in the trichoviruses, these two ORFs are slightly overlapped [15, 61, 62]. DaYuV1 is distantly related with AgVA, and their replicase and CP nt sequence similarity is below the 72% threshold value to demarcate species in the family *Betaflexiviridae* [15]. DaYuV1 and AgVA Rep and CP proteins were phylogenetically related in a distinct clade from other viruses belonging to genera within the subfamily *Trivirinae*. Thus, based on sequence similarities, genomic organization and phylogenetic relationships, DaYuV1 and AgVA should be classified as novel members of a new genus within the subfamily *Trivirinae* in the family *Betaflexiviridae*, which could be tentatively named as Daisagavirus.

The RhoDeV2 genome has three ORFs, with a genomic organization 5’-Rep-P2-P3-3’, that is similar to the one reported for the carrot chordoviruses [63]. Its replicase is distantly related with the one encoded by other betaflexiviruses, and the nt sequence similarity is below the 45% threshold value to demarcate genus in the family *Betaflexiviridae* [15]. Neither hits were obtained nor conserved domains were identified when the proteins encoded by ORFs 2 and 3 were analyzed, but a bipartite NLS was detected in the sequence of the ORF 3 encoded protein. Thus, RhoDeV2 ORF2 and ORF3 encoded proteins have an unknown function and are not likely a MP and CP, which are the putative proteins encoded by carrot chordoviruses ORFs2 and 3 [63], which likely indicate that RhoDeV2 is unique among betaflexiviruses. The RhoDeV2 replicase formed a monophyletic cluster that was related with the *Chordovirus* and *Ravavirus* clades. Thus, sequence similarities, the evolutionary relationships with other betaflexiviruses and the unknown function of its putative P2 and P3 proteins suggest that RhoDeV2 is a unique novel betaflexivirus that should belong to a novel genus within the subfamily *Trivirinae* in the family *Betaflexiviridae*, which could be tentatively named as Rhodovirus.

#### Deltaflexiviridae

The ATV3 and SesV1 genomic organization5’Rep-P2-P3-P4-P5-3’, resembles the one described for the deltaflexiviruses FgDFV1 [18]. Their replicase is distantly related with the one encoded by other deltaflexiviruses, and phylogenetic analysis showed that ATV3 and SesV1 grouped together with the deltaflexiviruses. Thus, based on sequence similarities, genomic organization and phylogenetic relationships, ATV3 and SesV1 should be classified as novel members of the genus *Deltaflexivrus* in the family *Deltaxiviridae*.

### Recombination

Interestingly, despite that the recombination is a key process that strongly impact the evolution of many plant virus species [26] and has been mentioned as a driving force in the evolution of flexiviruses [64], no recombination event, using as threshold a p-value < 0.05 in at least three algorithms, was detected in the alphaflexiviruses, betaflexiviruses and deltaflexiviruses identified in this study.

#### Tymoviridae and tymo-like virus

DaInvV1 and DaInv2 genome organization resemble the one described for marafiviruses [65], and these viruses are phylogenetically related with other marafiviruses. The highest aa identity when their CP was compared with other marafiviruses was below 80%, which is the criteria to demarcate species in the *Tymoviridae* family [21].Thus, based on sequence similarities, genomic organization and phylogenetic relationships, DaInvV1 and DaInv2 should be classified as novel members of the genus *Marafivirus* in the family *Tymoviridae*.

YePoV1 genomic organization 5’Rep-CP-3’, resembles the one reported for the tymovirids PYSaV [66] and GaTLV [67]. YePoV1 grouped together with these two viruses in the phylogenetic tree, and their CP shared an aa identity below 80%, which is the threshold value to demarcate species in the *Tymoviridae* family [21]. Thus, based on sequence similarities, genomic organization and phylogenetic relationships, YePoV1 would be the third member of the recently proposed genus *Gratylivirus*, that belongs to an unassigned family within the order *Tymovirales* [66, 67].

WaCraV1 has three ORFs and its genomic organization 5’-Rep-CP-P3-3’, is similar to the one reported for the invertebrate-associated macula-like viruses BeeMLV2 [68] and Bombyx mori Macula like-virus (BmMLV) [69], which differs from the one reported for the plant-associated maculaviruses [70, 71]. Interestingly, the P3 encoded by the WaCraV1 genome has a transmembrane domain, while no transmembrane domains were identified in the P3 encoded by BeeMLV2 and BmMLV genomes; thus the function of WaCraV1 P3 protein could be different than the BeeMLV2 and BmMLV P3s. WaCraV1 grouped together with BeeMLV2 and BmMLV in the phylogenetic tree, but their CP shared an aa identity lower than 80%, which is the criteria to demarcate species in the *Tymoviridae* family [21]. Phylogenetic relationships suggest that WaCraV1 is likely an invertebrate-infecting virus derived from insect RNA that was eventually co-purified with the targeted plant RNA. The identification of invertebrate-infecting viruses from plant transcriptome datasets was already reported elsewhere suggesting that some undetected invertebrate was contaminating the plant samples [72]. Based on sequence similarities, the distinct genomic organization and phylogenetic relationships WaCraV1, BeeMLV2 and BmMLV should be classified as members of a novel genus within the *Tymoviridae* family, which could be tentatively named as Inmaculavirus.

The KaV1 and PolWhV1 genome has three ORFs, 5’-Rep-P2-P3-3,’ where ORFs 1 and 2 are separated by a short intergenic region, while ORF3 is slightly overlapped with ORF2, which is likely a unique genomic organization among tymoviruses. KaV1 and PolWhV1 replicases are distantly related with other unclassified tymoviruses and mycotymoviruses, respectively, and the nt sequence similarity of its nearly complete genome is below the 80% threshold value to demarcate species in the family *Tymoviridae* [21]. No hits were obtained when the proteins encoded by ORFs 2 and 3 of KaV1 and PolWhV1 were analyzed, but a transmembrane domain was identified in the KaV1 encoded P2. Thus, KaV1 and PolWhV1 P2 and P3 have an unknown function and is tempting to speculate that KaV1 P2 could have a membrane associated function. Despite that Kav1 and PolWhV1 share a distinctive genomic organization among tymoviruses, their evolutionary history is different because the KaV1 replicase virus formed a monophyletic cluster that was distantly related with other tymoviruses, while PolWhV1 replicase clustered together with mycotymoviruses. Thus, the evolutionary relationship of KaV1 with other known members of the family *Tymoviridae*, and the unknown function of its putative P2 and P3 proteins suggest that KaV1 likely represents a novel genus in the *Tymoxiviridae* family which could be tentatively as Kavavirus. On the other hand, the evolutionary history of PolWhV1 suggest that this virus should be classified as a novel member of the proposed genus *Mycotymovirus* [23], despite its genomic organization is different than the reported mycotymoviruses [23, 73].

### Fungal RNA present in plant transcriptome datasets

It is tempting to speculate that fungal RNA was also present in the *Agave tequilana, Sesamum indicum* and *Triticum polonicum* transcriptomes given that the evolutionary analyses of ATV3, SesV1 and PolWhV1 suggest they couldbe derived from a fungus associated to these plant samples. Furthermore, an isolate of the fungi-associated deltaflexiviruses Erysiphe necator associated deltaflexivirus 2 and Alternaria alternate deltaflexivirus 1 were identified in the transcriptome of *Agrostis stolonifera* (MW328744). Recently the putative gammaflexivirus pistacia-associated flexivirus 1 was identified analysing a publically available transcriptome dataset of pistacia [39]. Thus, the identification of mycoviruses from plant transcriptome dataset its plausible and thus tentative host assignment should consider the potential impact of eventual mixed sampling of transcriptome datasets..

### Conclusion remarks

In summary, this study shows the complexity and diversity of the genome organization of viruses belonging to the order *Tymovirales* and reveal that analyzing SRA public data is a valuable tool not only to speed up the identification of novel viruses but also to increase our understanding of their evolution and taxonomical classification. Nevertheless, our inability to go back to the biological material which was employed to generate the transcriptome datasets analyzed, in order to check the viral genome sequences assembled and link the virus to a specific host, is the main limitation of using the data mining as a tool for virus discovery. This constraint may lead to a potential misidentification of the host species linked to the virus. Therefore a cautious attitude must be carried out before making strong statements related to viruses discovered from the analysis of transcriptome datasets publically available. This study includes the identification and characterization of a diverse set of tymovirids, presenting striking genome organization flexibility and interesting phylogenetic cues which expands the evolutionary history of the order incorporating 31 new tentative members of species and genera within the order, highlighting the importance of HTS in virus discovery and its potential in virus taxonomy.

## Acknowledgments

We would like to express a sincere gratitude to the generators of the underlying data used for this work, which are cited in Table 1. By following open access practices and supporting accessible raw sequence data in public repositories available to the research community, they have promoted the generation of new knowledge and ideas.

## Ethics declaration

### Human and animal rights statement

This manuscript presents original work that does not involve studies with humans or animals.

