## Supplementary material for "Exploring the tymovirids landscape through metatranscriptomics data": Supp Material: Supp Material.docx

**Figure S1**. A Maximum Likelihood phylogenetic based on a multiple amino acid alignment of the CP sequences of alphaflexiviruses, was constructed with the WAG+G+F model. Bootstrap values following 1000 replicates are given at the nodes, but only values above 40% are shown. Helenium virus S and Chrysanthemum virus B CP aa sequences were used as outgroup. The scale bar indicates the number of substitutions per site. Viruses identified in this study are noted with a green square. Accession numbers of every virus used to construct the ML tree are listed in Table S1.

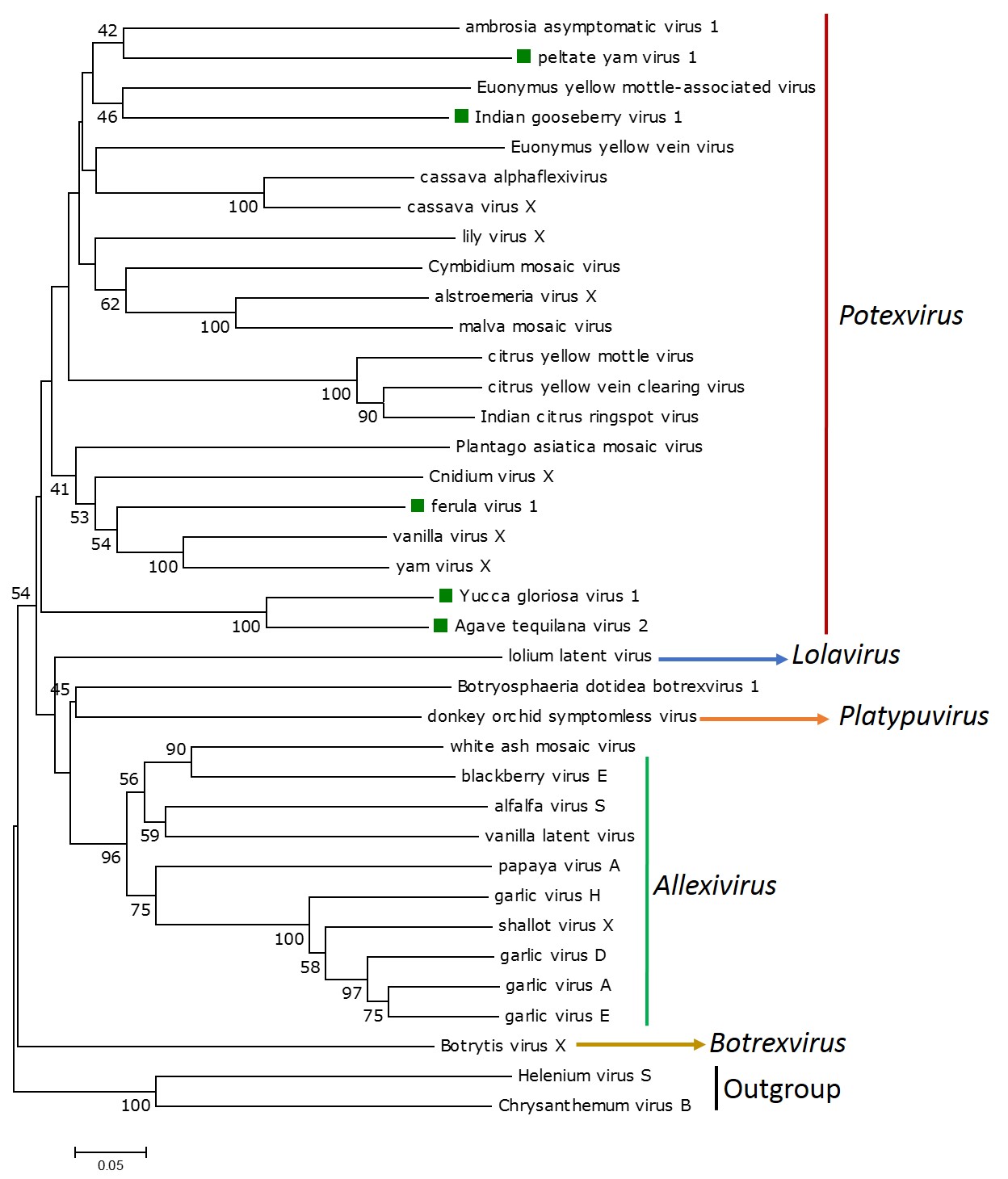

**Figure S2**. A Maximum Likelihood phylogenetic based on a multiple amino acid alignment of the CP sequences of viruses of the subfamily *Quinvirinae*, was constructed with the LG+G+F model. Bootstrap values following 1000 replicates are given at the nodes, but only values above 40% are shown. Apricot pseudo-chlorotic leaf spot virus, cherry mottle leaf virus and Grapevine Berry Inner necrosis virus CP aa sequences were used as outgroup. The scale bar indicates the number of substitutions per site. Viruses identified in this study are noted with a green square. Viruses identified with an * correspond to unpublished sequences derived from the analysis of transcriptome data publically available at the NCBI. Accession numbers of every virus used to construct the ML tree are listed in Table S1.

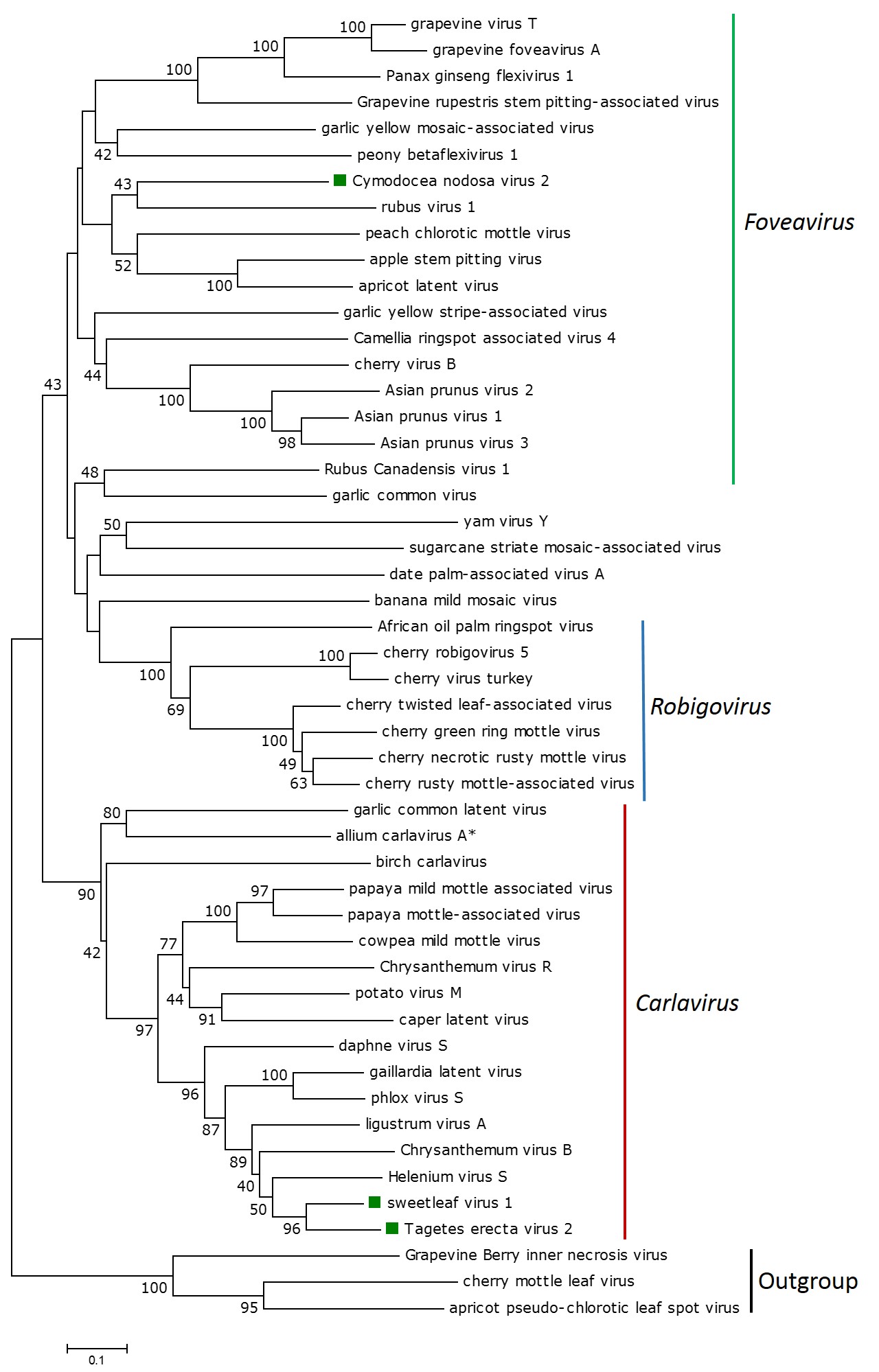

**Figure S3**. A Maximum Likelihood phylogenetic based on a multiple amino acid alignment of the CP sequences of viruses of the subfamily *Trinvirinae*, was constructed with the LG+G+F model. Bootstrap values following 1000 replicates are given at the nodes, but only values above 40% are shown. The scale bar indicates the number of substitutions per site. Viruses identified in this study are noted with a green square. Viruses identified with an * correspond to unpublished sequences derived from the analysis of transcriptome data publically available at the NCBI. Accession numbers of every virus used to construct the ML tree are listed in Table S1.

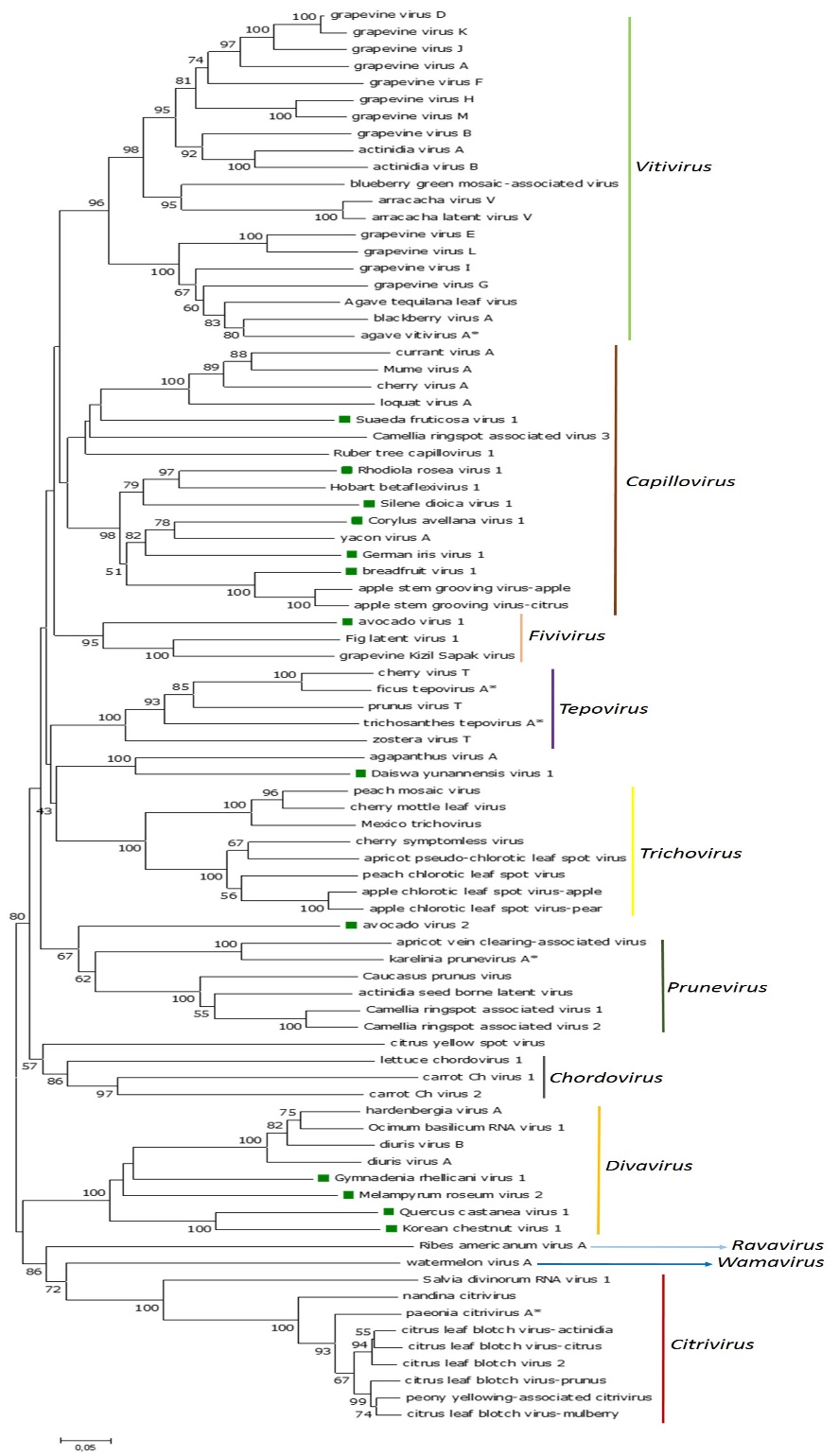

**Table S1**. Virus names, and NCBI accession numbers of tymovirids sequences used in this study

| Virus name | Accession number |
| --- | --- |
| actinidia virus A | LC491285 |
| actinidia virus B | JN427015 |
| actinidia seed borne latent virus | MF440375 |
| African oil palm ringspot virus | AY072921 |
| agapanthus virus A | MT533609 |
| Agave tequilana leaf virus | KY190215 |
| agave virus T | MW323519 |
| agave vitivirus A | MH898468 |
| alfalfa virus S | MN864567 |
| allium carlavirus A | MH898470 |
| alstroemeria virus X | AB206396 |
| Alternatia alternata deltaflexivirus 1 | MW567960 |
| ambrosia asymptomatic virus 1 | KF421905 |
| Anagyris vein yellowing virus | AY751780 |
| andean potato latent virus | JX508921 |
| apple chlorotic leaf spot virus-apple | LC533837 |
| apple chlorotic leaf spot virus-pear | KC935956 |
| apple stem grooving virus-apple | KU947036 |
| apple stem grooving virus-citrus | LC184612 |
| apple stem pitting virus | KF321966 |
| apricot latent virus | LC522982 |
| apricot pseudo-chlorotic leaf spot virus | AY713379 |
| apricot vein clearing-associated virus | KY132099 |
| arracacha latent virus V | KY451036 |
| arracacha virus V | KY392781 |
| Asian prunus virus 1 | KR998046 |
| Asian prunus virus 2 | KY310581 |
| Asian prunus virus 3 | KT893296 |
| asclepias asymtptomatic virus | HQ425778 |
| banana mild mosaic virus | AF314662 |
| bee macula-like virus 2 | MF998084 |
| birch carlavirus | MH536506 |
| blackberry virus A | MG254193 |
| blackberry virus E | JN053266 |
| blackberry virus S | FJ915122 |
| blueberry green mosaic-associated virus | MK460433 |
| Bombyx mori macula-like virus | KJ433990 |
| Botryosphaeria dotidea botrexvirus 1 | MT103578 |
| Botrytis cinerea mycotymovirus 1 | MN954873 |
| Botrytis virus F | MH338171 |
| Botrytis virus X | AY055762 |
| cactus virus X | JF937699 |
| Camellia-associated marafivirus | MT036048 |
| Camellia ringspot-associated virus 1 | MK050792 |
| Camellia ringspot-associated virus 2 | MK050793 |
| Camellia ringspot-associated virus 3 | MK050796 |
| Camellia ringspot-associated virus 4 | MT028514 |
| caper latent virus | MT311966 |
| carrot ch virus 1 | KF533711 |
| carrot ch virus 2 | KF533710 |
| cassava alphaflexivirus | KC505252 |
| cassava virus X | KY288487 |
| Caucasus prunus virus | KM507061 |
| chayote mosaic virus | AF195000 |
| cherry virus A | KY510919 |
| cherry virus B | LC373513 |
| cherry green ring mottle virus | KY178276 |
| cherry mottle leaf virus | AF170028 |
| cherry necrotic rusty mottle virus | LC522996 |
| cherry robigovirus 5 | MK035727 |
| cherry rusty mottle-associated virus | KF030870 |
| cherry symptomless virus | MK770441 |
| cherry twisted leaf-associated virus | KF030859 |
| cherry virus T | MT090966 |
| cherry virus turkey | MK600387 |
| Chrysanthemum virus B | AB245142 |
| Chrysanthemum virus R | MG432107 |
| citrus yellow spot virus | MN879752 |
| citrus leaf blotch virus-actinidia | MG604237 |
| citrus leaf blotch virus-citrus | MT078932 |
| citrus leaf blotch virus-mulberry | MT767171 |
| citrus leaf blotch virus-prunus | KR023647 |
| citrus leaf blotch virus 2 | MH558590 |
| citrus sudden death-associated virus | KY110736 |
| citrus virus C | MN879754 |
| citrus yellow mottle virus | MK957246 |
| citrus yellow vein clearing virus | KT696510 |
| Cnidium virus X | LC460456 |
| cowpea mild mottle virus | MW534944 |
| currant virus A | KT763043 |
| Cymbidium mosaic virus | U62963 |
| daphne virus S | AJ620300 |
| date palm-associated virus A | BK011997 |
| Diascia yellow mottle virus | EU684141 |
| diuris virus A | JX173276 |
| diuris virus B | JX133277 |
| donkey orchid symptomless virus | KC923234 |
| dulcamara mottle virus | AY789137 |
| eggplant mosaic virus | KJ690172 |
| Entoleuca gammaflexivirus 1 | MF375883 |
| Entoleuca gammaflexivirus 2 | MF375884 |
| Eryshipe necator associated deltaflexivirus 1 | MN627486 |
| Eryshipe necator associated deltaflexivirus 2 | MN627444 |
| Eryshipe necator associated deltaflexivirus 3 | MN627465 |
| Eryshipe necator associated deltaflexivirus 4 | MN627471 |
| Euonymus yellow mottle-associated virus | MK572000 |
| Euonymus yellow vein virus | MF078061 |
| ficus tepovirus A | MH898491 |
| fig latent virus 1 | FN377573 |
| Fusarium graminearum deltaflexivirus 1 | KX015962 |
| Fusarium graminearum mycotymovirus 1 | KT360947 |
| gaillardia latent virus | KJ415259 |
| garlic common latent virus | JF320810 |
| garlic common virus | MN059101 |
| garlic virus A | MT731489 |
| garlic virus D | MN059343 |
| garlic virus E | AJ292230 |
| garlic virus H | MN059332 |
| garlic yellow mosaic-associated virus | MH120170 |
| garlic yellow stripe-associated virus | MT981417 |
| Glehnia littoralis marafivirus | BK013331 |
| grapevine-associated tymo-like virus | MH383239 |
| grapevine berry inner necrosis virus | KU971246 |
| grapevine foveavirus A | MN553040 |
| grapevine fleck virus | AJ309022 |
| grapevine Kizil Sapak virus | MN172165 |
| grapevine Pinot gris virus | MN458445 |
| grapevine red glove virus | KX109927 |
| Grapevine rupestris stem pitting-associated virus | JQ922417 |
| Grapevine rupestris vein feathering virus | MN974274 |
| grapevine Syrah virus 1 | FJ436028 |
| grapevine virus A | DQ855082 |
| grapevine virus B | MN716773 |
| grapevine virus D | MF774336 |
| grapevine virus E | MF991950 |
| grapevine virus F | LC617946 |
| grapevine virus G | MF405923 |
| grapevine virus H | MN716768 |
| grapevine virus I | MF927925 |
| grapevine virus J | MG637048 |
| grapevine virus K | MF072319 |
| grapevine virus L | MT269788 |
| grapevine virus M | MK492703 |
| grapevine virus Q | FJ977041 |
| grapevine virus T | MG195287 |
| hardenbergia virus A | HQ241409 |
| helenium virus S | MW207172 |
| Hobart betaflexivirus 1 | MG995738 |
| hosta virus X | AJ620114 |
| Hypera postica associated alphaflexivirus | MW676130 |
| Indian citrus ringspot virus | AF406744 |
| insect-associated alphaflexivirus 1 | MN203143 |
| karelinia prunevirus A | MH898497 |
| Kennedya yellow mosaic virus | DD00637 |
| lettuce chordovirus 1 | MG208123 |
| Lentinula edodes deltaflexivirus 1 | MN744723 |
| Lentinula edodes deltaflexivirus 2 | MN744724 |
| ligustrum virus A | MN786957 |
| lily virus X | LC335818 |
| lolium latent virus | EU489641 |
| loquat virus A | MK936045 |
| malva mosaic virus | DQ660333 |
| maize rayado fino virus | AF265566 |
| Medicago sativa marafivirus 1 | MF443260 |
| Mexico trichovirus | MK012336 |
| mume virus A | MG783575 |
| nandina citrivirus | MN055483 |
| naranjilla chlorotic mosaic virus | MG323924 |
| naranjilla mild mosaic virus | MH784952 |
| nectarine virus M | KT273413 |
| nemesia ring necrosis virus | AY751778 |
| oat blue dwarf virus | GU396990 |
| Ocimum basilicum RNA virus 1 | MF196913 |
| olive latent virus 3 | FJ444852 |
| okra mosaic virus | EF554577 |
| paeonia citrivirus A | MH898501 |
| Panax ginseng flexivirus 1 | MH036372 |
| papaya mild mottle-associated virus | MK984598 |
| papaya mottle-associated virus | MK984600 |
| papaya virus A | MN418120 |
| passion fruit yellow mosaic virus | KY823429 |
| peach chlorotic leaf spot virus | MH084695 |
| peach chlorotic mottle virus | EF693898 |
| peach virus D | MH898957 |
| peony betaflexivirus 1 | MN253489 |
| peony yellowing-associated citrivirus | MN253488 |
| Pestalotiopsis deltaflexivirus 1 | MW017460 |
| phlox virus S | EF492068 |
| Physalis mottle virus | Y16104 |
| pistacia-associated flexivirus 1 | MK605686 |
| Plantago asiatica mosaic virus | KU697313 |
| plantago mottle virus | AY751778 |
| Poinsettia mosaic virus | AM412237 |
| potato virus M | D14449 |
| potato virus T | AB697482 |
| prunus virus T | KF700262 |
| prunus yellow spot-associated virus | MK404133 |
| Ribes americanum virus A | MF166685 |
| ruber tree capillovirus 1 | MN047299 |
| Rubus canandensis virus 1 | JX277553 |
| rubus virus 1 | MN944023 |
| Salvia divinorum RNA virus 1 | MH899178 |
| Sclerotinia sclerotiorum debilitation-associated RNA virus 1 | AY147260 |
| Sclerotinia sclerotiorum debilitation-associated RNA virus 2 | KJ561219 |
| Sclerotinia sclerotiorum deltaflexivirus 2 | MH299810 |
| Sclerotinia sclerotiorum mycotymovirus 1 | MK530705 |
| Sclerotinia sclerotiorum mycotymovirus 2 | MT706023 |
| Scrophularia mottle virus | AY751777 |
| sorghum bicolor marafivirus | MN100128 |
| shallot virus X | MH389250 |
| soybean-leaf associated mycoflexivirus 1 | KT598226 |
| strawberry mild yellow edge virus | KR350470 |
| sugarcane striate mosaic-associated virus | AF315308 |
| switchgrass mosaic virus | JF727261 |
| tomato yellow blotch virus | EU779803 |
| trichosanthes tepovirus A | MH898525 |
| turnip yellow mosaic virus | KJ690173 |
| ullucus tymovirus 1 | MH645153 |
| ullucus tymovirus 2 | MH645152 |
| vanilla latent virus | MF150239 |
| vanilla virus X | MF150240 |
| watercress white vein virus | JQ001816 |
| watermelon virus A | KY363796 |
| white ash mosaic virus | DQ412998 |
| yacon virus A | KU375548 |
| yam virus X | KJ711908 |
| yam virus Y | MK782910 |
| zostera virus T | MK514426 |

**Table S2**. Summary of assembly statistics of the tymovirids sequences identified from the transcriptome data available in the NCBI database.

| **Virus name** | **Abbreviation** | **Bioproject ID** | **Total virus reads** | **Mean Coverage** | **Reads per Millon** |
| --- | --- | --- | --- | --- | --- |
| *Agave tequilana* virus 2 | ATV1 | PRJNA193469 | **7300** | **167.1X** | **421.9** |
| *Agave tequilana* virus 3 | ATV3 | PRJNA193469 | **5927** | **103.8X** | **307.1** |
| Avocado virus 1 | AvoV1 | PRJNA391003 | **9450** | **175.9X** | **45.7** |
| Avocado virus 2 | AvoV2 | PRJNA391003 | **8547** | **155.5X** | **32.9** |
| breadfruit virus 1 | BreFV1 | PRJNA311339 | **6570** | **151.6X** | **305.6** |
| *Corylus avellana* virus 1 | CorAvV1 | PRJNA316492 | **7325** | **170.9X** | **327.1** |
| *Cymodocea nodosa* virus 1 | CyNoV1 | PRJNA275569 | **5548** | **151.1X** | **226.4** |
| *Cymodocea nodosa* virus 2 | CyNoV2 | PRJNA275569 | **4841** | **115.4X** | **192.8** |
| *Daiswa yunannensis* virus 1 | DaYuV1 | PRJNA352766 | **6078** | **138.3X** | **205.3** |
| *Davidia involucrata* virus 1 | DaInvV1 | PRJNA513477 | **5163** | **108.9X** | **230.5** |
| *Davidia involucrata* virus 2 | DaInvV2 | PRJNA513477 | **4933** | **109.2X** | **220.2** |
| Ferula virus 1 | FerV1 | PRJNA476150 | **6860** | **174.6X** | **181.5** |
| German iris virus 1 | GerIV1 | PRJNA638517 | **2953** | **83.8X** | **45.4** |
| *Gymnadenia rhellicani* virus 1 | GymRhV1 | PRJNA504609 | **8120** | **162.6X** | **416.4** |
| Indian gooseberry virus 1 | InGoV1 | PRJNA431341 | **8407** | **201.1X** | **230.3** |
| Kava virus 1 | KaV1 | PRJNA494686 | **3535** | **55.87X** | **20.5** |
| Korean chestnut virus 1 | KoChV1 | PRJNA534190 | **5173** | **105.6X** | **145.7** |
| *Melampyrum roseum* virus 1 | MelRoV2 | PRJDB5395 | **6765** | **143.1X** | **276.1** |
| Peltate yam virus 1 | PelYV1 | PRJNA243000 | **5209** | **124.9X** | **132.2** |
| Polish wheat virus 1 | PolWhV1 | PRJNA304266 | **4945** | **96.3X** | **42.15** |
| *Quercus castanea* virus 1 | QueCaV1 | PRJNA532454 | **5632** | **108.8X** | **264.4** |
| *Rhodiola rosea* virus 1 | RhRoV1 | PRJNA398393 | **4053** | **97.11X** | **115.8** |
| *Rhododendron delavayi* virus 2 | RhoDeV2 | PRJNA358123 | **3132** | **60.5X** | **79.1** |
| Sesame virus 1 | SesV1 | PRJNA74261 | **6527** | **117.1X** | **287.4** |
| *Silene dioica* virus 1 | SiDiV1 | PRJNA356362 | **6413** | **136.7X** | **393.4** |
| *Suaeda fruticosa* virus 1 | SuFruV1 | PRJNA279892 | **4274** | **96.56X** | **247.1** |
| Sweetleaf virus 1 | SleV1 | PRJNA591974 | **8140** | **147.6X** | **317.9** |
| *Tagetes erecta* virus 2 | TaEV2 | PRJNA431782 | **9076** | **160.1X** | **444.9** |
| Watercress associated virus 1 | WaCraV1 | PRJNA284126 | **4166** | **92.6X** | **148.2** |
| yellow poplar virus 1 | YePoV1 | PRJNA556244 | **5384** | **133.1X** | **113.5** |
| *Yucca gloriosa* virus 1 | YuGlV1 | PRJNA374584 | **8456** | **200.1X** | **177.2** |

**Table S3**. Functional domains identified in the encoded protein by every tymovirid assembled in this study

| Virus name | Protein name | Conserved domains | Pfam | aa position |
| --- | --- | --- | --- | --- |
| *Agave tequilana* virus 2 | Replicase  Triple gene block 1  Triple gene block 2  Triple gene block 3  Capsid protein | Viral methyltransferase  Viral_helicase1  RdRP_2  Viral_helicase1  Plant viral movement protein  Triple gene block 3  Flexi_CP | PF01660  PF01443  PF00978  PF01443  PF01307  PF02495  PF00286 | 40-322  71-951  1103-1429  25-218  4-103  24-78  93-230 |
| *Agave tequilana* virus 3 | Replicase  Protein 2  Protein 3  Protein 4  Protein 5 | Viral methyltransferase  RdRP_2  -  -  -  - | PF01660  PF00978  -  -  -  - | 210-516  1750-2085  -  -  -  - |
| Avocado virus 1 | Replicase  Movement protein  Capsid protein  Protein 4  Nucleic acid binding protein | Viral methyltransferase  Viral_helicase1  RdRP_2  Viral movement protein  Trichovirus coat protein  -  Carlavirus putative NABP | PF01660  PF01443  PF00978  PF01107  PF05892  -  PF01623 | 44-334  985-1227  1388-1773  37-168  48-210  -  5-73 |
| Avocado virus 2 | Replicase  Movement protein  Capsid protein | Viral methyltransferase  2OG-FeII_Oxy_2  Viral_helicase1  RdRP_2  Viral movement protein  Trichovirus coat protein | PF01660  PF13532  PF01443  PF00978  PF01107  PF05892 | 44-348  884-1029  1227-1483  1623-2017  11-190  24-203 |
| Breadfruit virus 1 | Polyprotein  Movement protein | Viral methyltransferase  DUF1717  Viral_helicase1  RdRP_2  Trichovirus coat protein  Viral movement protein | PF01660  PF05414  PF01443  PF00978  PF05892  PF01107 | 43-347  635-712  791-1048  1221-1510  1891-2103  10-201 |
| *Corylus avellana* virus 1 | Polyprotein  Movement protein | Viral methyltransferase  DUF1717  Viral_helicase1  RdRP_2  Trichovirus coat protein  Viral movement protein | PF01660  PF05414  PF01443  PF00978  PF05892  PF01107 | 43-349  607-684  767-1019  1178-1480  1926-2087  10-203 |
| *Cymodocea nodosa* virus 1 | Replicase  Movement protein | Viral methyltransferase  Viral_helicase1  RdRP_2  3A/RNA2 MP family | PF01660  PF01443  PF00978  PF00803 | 38-321  656-890  1037-1336  12-239 |
| *Cymodocea nodosa* virus 2 | Replicase*  Triple gene block 1  Triple gene block 2  Triple gene block 3  Capsid protein | Viral_helicase1  RdRP_2  Viral_helicase1  Plant viral movement protein  Triple gene block 3  Flexi_CP | PF01443  PF00978  PF01443  PF01307  PF02495  PF00286 | 613-873  994-1407  22-221  4-104  6-64  39-176 |
| *Daiswa yunannensis* virus 1 | Replicase  Movement protein  Capsid protein | Viral methyltransferase  Viral_helicase1  RdRP_2  Viral movement protein  Trichovirus coat protein | PF01660  PF01443  PF00978  PF01107  PF05892 | 41-377  841-1084  1230-1578  11-190  24-203 |
| *Davidia involucrata* virus 1 | Polyprotein | Viral methyltransferase  Tymovirus endopeptidase  Viral_helicase1  RdRP_2  Tymovirus coat protein | PF01660  PF05381  PF01443  PF00978  PF00983 | 214-498  927-1034  1117-1350  1633-1921  2080-2262 |
| *Davidia involucrata* virus 2 | Polyprotein | Viral methyltransferase  Tymovirus endopeptidase  Viral_helicase1  RdRP_2  Tymovirus coat protein | PF01660  PF05381  PF01443  PF00978  PF00983 | 144-426  878-984  1064-1297  1547-1828  1991-2167 |
| Ferula virus 1 | Replicase  Triple gene block 1  Triple gene block 2  Triple gene block 3  Capsid protein | Viral methyltransferase  Viral_helicase1  RdRP_2  Viral_helicase1  Plant viral movement protein  Triple gene block 3  Flexi_CP | PF01660  PF01443  PF00978  PF01443  PF01307  PF02495  PF00286 | 39-325  576-809  969-1273  34-224  4-98  5-61  25-162 |
| German iris virus 1 | Polyprotein*  Movement protein | DUF1717  Viral_helicase1  RdRP_2  Trichovirus coat protein  Viral movement protein | PF05414  PF01443  PF00978  PF05892  PF01107 | 178-255  338-592  754-1085  1539-1724  10-203 |
| *Gymnadenia rhellicani* virus 1 | Polyprotein  Movement protein  Nucleic acid binding protein | Viral methyltransferase  Viral_helicase1  RdRP_2  Trichovirus coat protein  Viral movement protein  Viral_NABP | PF01660  PF01443  PF00978  PF05892  PF01107  PF05515 | 43-368  872-1133  1285-1688  2085-2227  42-197  47-97 |
| Indian gooseberry virus 1 | Replicase  Triple gene block 1  Triple gene block 2  Triple gene block 3  Capsid protein | Viral methyltransferase  Viral_helicase1  RdRP_2  Viral_helicase1  Plant viral movement protein  Triple gene block 3  Flexi_CP | PF01660  PF01443  PF00978  PF01443  PF01307  PF02495  PF00286 | 39-336  611-844  1001-1303  26-226  6-110  1-59  82-221 |
| Kava virus 1 | Replicase  Protein 2  Protein 3 | Viral methyltransferase  Viral_helicase1  RdRP_2  Transmembrane  - | PF01660  PF01443  PF00978  - | 81-378  1283-1519  1896-2165  7-29  - |
| Korean chestnut virus 1 | Polyprotein  Movement protein | Viral methyltransferase  Viral_helicase1  RdRP_2  Trichovirus coat protein  Viral movement protein | PF01660  PF01443  PF00978  PF05892  PF01107 | 44-350  931-1177  1322-1684  2145-2327  12-195 |
| *Melampyrum roseum* virus 2 | Polyprotein  Movement protein  Nucleic acid binding protein | Viral methyltransferase  Viral_helicase1  RdRP_2  Trichovirus coat protein  Viral movement protein  Viral_NABP | PF01660  PF01443  PF00978  PF05892  PF01107  PF05515 | 1-233  739-1001  1159-1500  1955-2117  11-194  65-115 |
| Peltate yam virus 1 | Replicase  Triple gene block 1  Triple gene block 2  Triple gene block 3  Capsid protein | Viral methyltransferase  Viral_helicase1  RdRP_2  Viral_helicase1  Plant viral movement protein  Triple gene block 3  Flexi_CP | PF01660  PF01443  PF00978  PF01443  PF01307  PF02495  PF00286 | 39-323  595-828  975-1277  25-230  4-103  20-80  42-182 |
| Polish wheat virus 1 | Replicase  Protein 2  Protein 3 | Viral methyltransferase  Tymovirus endopeptidase  Viral_helicase1  RdRP_2  -  - | PF01660  PF05381  PF01443  PF00978  -  - | 440-720  1142-1214  1351-1588  1741-2099  -  - |
| *Quercus castanea* virus 1 | Polyprotein  Movement protein | Viral methyltransferase  Viral_helicase1  RdRP_2  Trichovirus coat protein  Viral movement protein | PF01660  PF01443  PF00978  PF05892  PF01107 | 43-347  903-1165  1304-1670  2146-2328  28-196 |
| *Rhodiola rosea* virus 1 | Polyprotein  Movement protein | Viral methyltransferase  DUF1717  Viral_helicase1  RdRP_2  Trichovirus coat protein  Viral movement protein | PF01660  PF05414  PF01443  PF00978  PF05892  PF01107 | 43-349  571-631  732-997  1135-1462  1871-2048  14-201 |
| *Rhododendron delavayi* virus 2 | Replicase  Protein 2  Protein 3 | Viral methyltransferase  2OG-FeII_Oxy_2  Viral_helicase1  RdRP_2  -  - | PF01660  PF13532  PF01443  PF00978  -  - | 44-367  647-791  1085-1325  1449-1807  -  - |
| Sesame virus 1 | Replicase  Protein 2  Protein 3  Protein 4  Protein 5 | Viral methyltransferase  Viral_helicase1  RdRP_2  -  -  -  - | PF01660  PF01443  PF00978  -  -  -  - | 210-516  1233-1482  1750-2085  -  -  -  - |
| *Silene dioica* virus 1 | Polyprotein  Movement protein | Viral methyltransferase  DUF1717  Viral_helicase1  RdRP_2  Trichovirus coat protein  Viral movement protein | PF01660  PF05414  PF01443  PF00978  PF05892  PF01107 | 43-349  592-635  739-1003  1143-1482  2070-2257  12-201 |
| *Suaeda fruticosa* virus 1 | Polyprotein  Movement protein | Viral methyltransferase  Viral_helicase1  RdRP_2  Trichovirus coat protein  Viral movement protein | PF01660  PF01443  PF00978  PF05892  PF01107 | 43-347  720-967  1096-1483  2025-2209  16-203 |
| Sweetleaf virus 1 | Replicase  Triple gene block 1  Triple gene block 2  Triple gene block 3  Capsid protein  Nucleic acid binding protein | Viral methyltransferase  Carlavirus endopeptidase  Viral_helicase1  RdRP_2  Viral_helicase1  Plant viral movement protein  Triple gene block 3  Carlavirus coat  Flexi_CP  Carlavirus putative NABP | PF01660  PF05379  PF01443  PF00978  PF01443  PF01307  PF02495  PF08358  PF00286  PF01623 | 43-354  982-1066  1153-1412  1529-1946  24-223  4-103  4-60  47-98  107-246  7-96 |
| *Tagetes erecta* virus 2 | Replicase  Triple gene block 1  Triple gene block 2  Triple gene block 3  Capsid protein  Nucleic acid binding protein | Viral methyltransferase  Carlavirus endopeptidase  Viral_helicase1  RdRP_2  Viral_helicase1  Plant viral movement protein  Triple gene block 3  Carlavirus coat  Flexi_CP  Carlavirus putative NABP | PF01660  PF05379  PF01443  PF00978  PF01443  PF01307  PF02495  PF08358  PF00286  PF01623 | 43-354  983-1069  1155-1415  1540-1949  24-223  4-103  4-60  48-99  108-247  6-97 |
| Watercress associated virus 1 | Replicase  Coat Protein  Protein 3 | Viral methyltransferase  Tymovirus endopeptidase  Viral_helicase1  RdRP_2  Tymovirus coat protein  Transmembrane | PF01660  PF05381  PF01443  PF00978  PF00983 | 38-319  718-825  905-1139  1413-1720  90-270  17-39 |
| yellow poplar virus 1 | Replicase  Protein 2 | Viral methyltransferase  Viral_helicse1  RdRP_2  - | PF01660  PF01443  PF00978  - | 92-395  956-1190  1377-1674  - |
| *Yucca gloriosa* virus 1 | Replicase  Triple gene block 1  Triple gene block 2  Triple gene block 3  Capsid protein | Viral methyltransferase  Viral_helicase1  RdRP_2  Viral_helicase1  Plant viral movement protein  Triple gene block 3  Flexi_CP | PF01660  PF01443  PF00978  PF01443  PF01307  PF02495  PF00286 | 40-322  659-892  1044-1313  25-218  4-103  5-62  96-233 |
